# Targeted shedding of extracellular membrane proteins by induced protease recruitment

**DOI:** 10.64898/2026.02.17.706468

**Authors:** Zi Yao, Fangzhu Zhao, Kun Miao, Trenton M. Peters-Clarke, Yun Zhang, Snehal D. Ganjave, Angel L. Vázquez-Maldonado, Yan Wu, Kaan Kumru, Hammam Jumaa, Kevin K. Leung, James A. Wells

## Abstract

Extracellular targeted protein degradation has emerged as a promising therapeutic modality to eliminate proteins of interest (POIs) at the cell surface, by using bifunctional molecules to recruit natural recycling receptors or membrane-bound E3 ligases that redirect POIs to the lysosome. Another natural mechanism involves extracellular proteases that cleave and shed extracellular domains. Here, we exploit this endogenous mechanism by engineering bispecific antibody *Shedders*, that recruit a classic sheddase ADAM10 to POIs, inducing selective ectodomain shedding. We first targeted the immune checkpoint receptor LAG-3 and observed robust depletion of surface LAG-3 accompanied by accumulation of soluble LAG-3 fragments in both engineered cell lines and primary human T cells. Using biochemical and imaging assays, we confirmed that this antibody-induced shedding is restricted to extracellular protease activity and occurs independently of lysosomal trafficking. Notably, induced shedding of LAG-3 on activated primary T cells partially alleviated inhibitory signaling and reinvigorated IFN*γ* secretion. We extended the scope of induced shedding by developing *Shedders* that recognize synthetic epitope-tags that enabling rapid assessment of substrate compatibility across diverse targets. Using this platform, we identified multiple immune modulatory cell-surface receptors, including IL6Rα, CD62L and MIC-A that can be targeted for shedding. In summary, this work establishes a new paradigm for targeted extracellular proteolysis and expands the toolkit for studying extracellular proteolysis with potential therapeutic benefit.

## INTRODUCTION

Targeted protein degradation (TPD) is a transformative strategy to target “undruggable” proteins through induced proximity^1,2^. Bifunctional molecules are engineered to recruit a protein of interest (POI) to the cell’s endogenous degradation machinery^1^. Historically, TPD platforms have focused on recruiting cytosolic E3 ligases to induce the ubiquitination of intracellular proteins, thereby promoting their subsequent degradation via the proteasome^3^. Degrading disease-driving POI offers several distinct advantages over traditional small-molecule inhibition, including the elimination of scaffolding functions and the ability to overcome clinical resistance mutations^3,4^. More recently, this proximity-based strategy has been extended to the extracellular proteome by trafficking cell-surface POIs to the lysosome for degradation^2,5,6^.

A myriad of extracellular TPD (eTPD) strategies have been developed by hijacking endogenous glycoproteins^5^, neonatal Fc receptors^7^, membrane-tethered E3 ligases^6,8^, and various metabolic or recycling receptors^9,10^ for internalization and proteolysis in the lysosome **(Figure 1a)**. These platforms have demonstrated remarkable success in eliminating oncogenic signaling receptors, immune checkpoint proteins, and soluble growth factors. Despite the broad scope, all existing eTPD platforms rely on internalization machinery to shuttle POIs to the lysosome for complete degradation^2,5,7^. Another major role of cellular proteolysis is by extracellular proteases that become activated at the cell surface and lead to nicking for site-specific shedding of extracellular domains. Recently many of these events have been annotated by N-terminomics of the residual fragments either shed or remaining on the cell surface. The role of most of these events has not been annotated but many of these site-specific proteolysis events are gain-of-function events such as shedding of growth factors and receptors.

**Figure 1.**
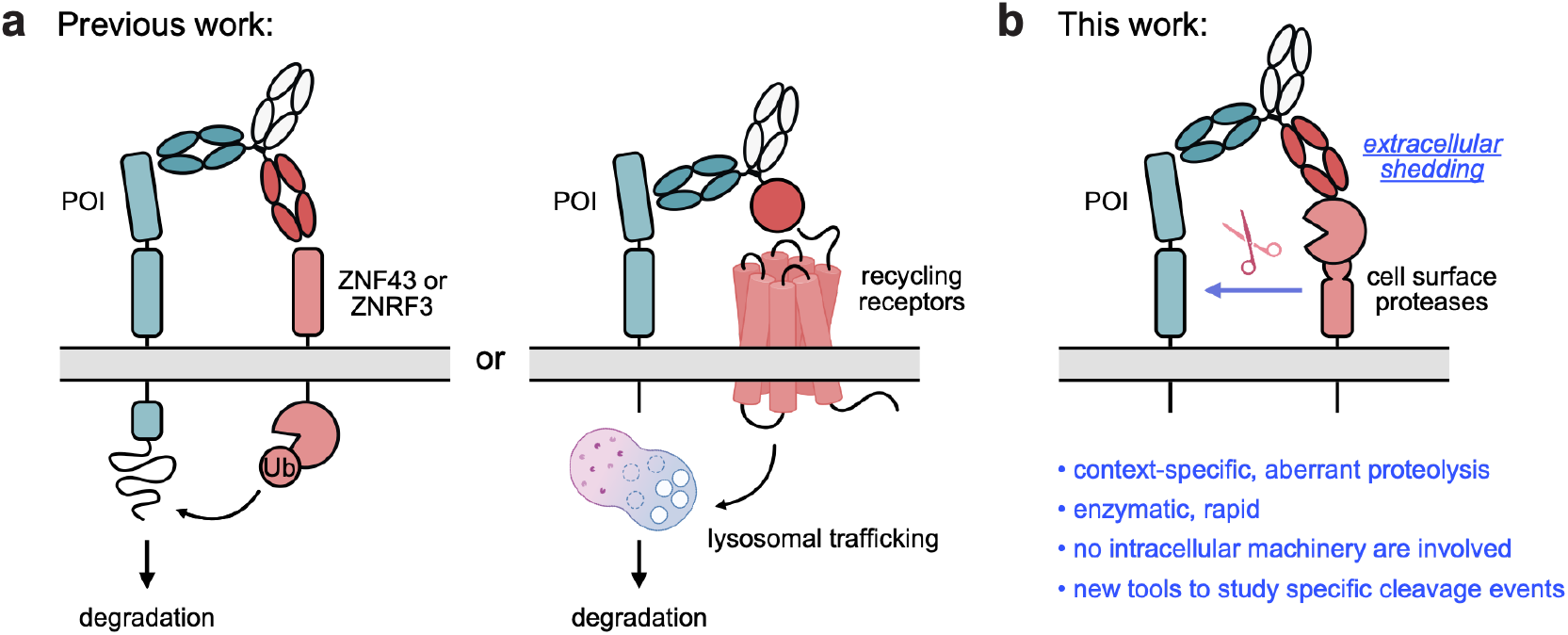
Induced proteolysis via bispecific antibodies. **(a)** Current strategies for extracellular targeted protein degradation. Bispecific antibodies are designed to simultaneously bind a protein of interest (POI) and either a membrane-tethered E3 ligase (*left*) or a recycling receptor (*right*). Upon complex formation, the POI is internalized and degraded in the lysosome. **(b)** Induced proteolysis via protease recruitment. Instead of engaging a recycling receptor, bispecific shedders recruit membrane-bound proteases to the POI, triggering its cleavage and release from the cell surface.

Here, we envision a new paradigm for studying and inducing extracellular proteolysis by engineering bispecific antibodies that recruit extracellular proteases to a POI via induced ectodomain shedding **(Figure 1b)**. Unlike existing modalities for eTPD, surface proteolysis takes place entirely outside of the cell. Furthermore, because this mechanism involves a direct enzymatic reaction rather than multi-step trafficking, it is likely more kinetically favorable and achieves catalytic degradation. Extracellular proteases and sheddases are ubiquitous on the cell surface and play a vital role in regulating transmembrane proteins^11,12^. By cleaving and releasing ectodomains, these enzymes can alter the signaling behavior and function of their substrates^13– 15^. Among the key enzymes are the a disintegrin and metalloproteinase (ADAM) family members^16,17^, most notably ADAM10 and ADAM17^18^. These proteases possess broad substrate repertoires that span diverse physiological and pathological contexts^19,20^. Consequently, hijacking ADAM-family proteases for induced shedding offers a versatile platform to modulate disease biology and provides novel opportunities for therapeutic intervention.

Substrates of ADAM10 represent both physiological regulators and potential pathological drivers in contexts such as inflammation^21^, neurodegeneration^22^, and cancer^23,24^. We initially focus on one well-characterized natural substrate: lymphocyte activation gene 3 (LAG-3), an immune checkpoint receptor that plays a pivotal role in regulating T cell activation and antitumor immunity^25,26^. LAG-3 interacts with multiple ligands, including the major histocompatibility complex class II (MHC II) on antigen-presenting cells (APCs)^27^ and the fibrinogen-like protein 1 (FGL1)^28^ secreted by hepatocytes and certain tumor types. Its inhibitory signaling on T cells is finely regulated by proteolytic shedding mediated by ADAM10 and ADAM17^29^, which cleave LAG-3 in its membrane-proximal region to release the soluble LAG-3 (sLAG-3)^29^. The cleavage of sLAG-3 has been well-documented biochemically and in proteomic studies, yet sLAG-3 itself is not known to have any biological activity. Importantly, *in vitro* and *in vivo* studies have shown that blocking LAG3-shedding via engineered non-cleavable LAG-3 mutants exacerbate inhibitory phenotypes on T cells^29^. Mice expressing the non-cleavable LAG-3 mutant exhibit resistance to immunotherapy and fail to mount effective antitumor immune responses^30^, underscoring the biological relevance of regulated LAG-3 shedding in immune modulation.

Based on these insights, we engineer bispecific antibody shedders designed to recruit ADAM10 to LAG-3 via induced proximity, promoting potent and rapid extracellular shedding. This induced shedding strategy enhances T cell activation, demonstrating greater potency than the clinically approved LAG-3-blocking antibodies. Beyond LAG-3, we also show that multiple natural membrane substrates can be targeted for shedding via ADAM10 using this induced proximity approach. We further demonstrated broad utility by establishing a universal shedding assay for investigating specific proteolytic event. Overall, our work establishes a novel modality that harnesses the endogenous shedding machinery in cells to achieve targeted shedding of disease-relevant surface proteins. This strategy expands the toolkit for extracellular proteolysis and offers potential for novel biologic therapies.

## RESULTS

### ADAM10 recruitment induces LAG3 shedding

To induce proximity between ADAM10 and LAG-3, we engineered bispecific IgG constructs with silenced Fc effector function **(Figure 2a)**^10,31^. For the protease-recruitment arm, we selected the previously characterized clone 11G2^32^, which targets the disintegrin domain of ADAM10. 11G2 has been shown not to interfere with ADAM10 catalytic activity, making it an attractive binder for shedder design. For POI targeting arm, we co-opted the clinically approved antibody Relatlimab^33^, which recognizes the N-terminal Domain 1 (D1) of LAG-3. Relatlimab has a high affinity for LAG-3, which likely favors the formation of a stable ternary complex^33^. Furthermore, because D1 is distal to both the annotated cut site and the membrane-proximal region, we reasoned that targeting this epitope would minimize steric occlusion and enable flexibility to for the protease to access the cleavage site. For ease of expression, we formatted the LAG-3 binding arm as a single-chain variable fragment (scFv) while the ADAM10 binding arm was formatted as a scFv (LAG-3 shedder 1) or Fab (LAG-3 shedder 2).

**Figure 2.**
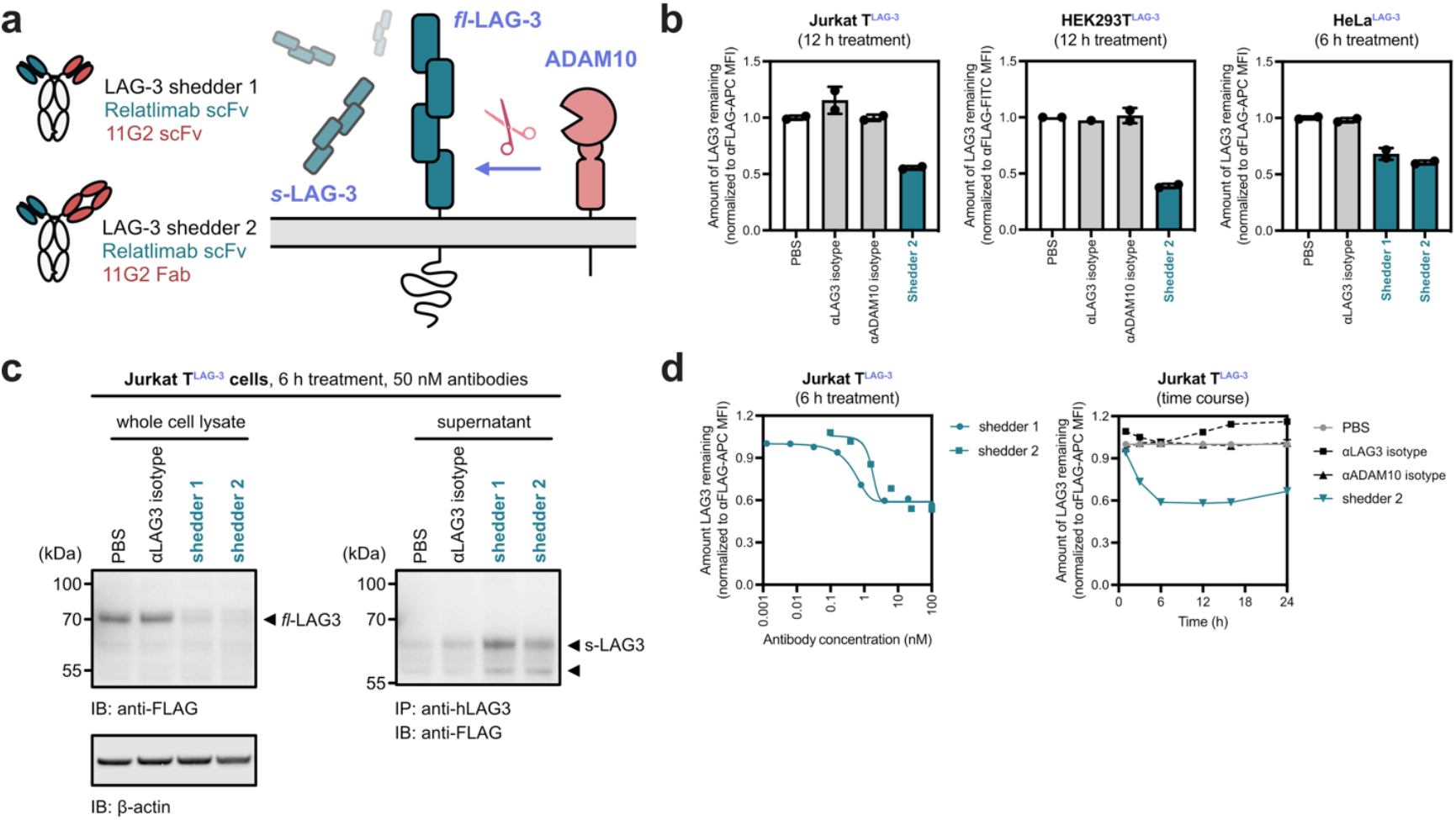
Induced shedding of LAG-3. **(a)** LAG-3 is endogenously cleaved by ADAM10, releasing a ∼60 kDa soluble fragment. Two bispecific shedders were designed to recruit ADAM10 to enhance the proteolysis of LAG-3. **(b)** Cells stably expressing LAG-3 were treated with bispecific shedders or control antibodies. Flow cytometry revealed robust depletion of surface LAG-3 in the presence of shedders. Percent LAG3 was determined by median fluorescence intensity (MFI) of the APC fluorescence channel of live cells. Each sample was tested in biological duplicate and error bars represented standard deviation. **(c)** Western blot analysis of Jurkat cells overexpressing LAG-3 showed loss of full-length LAG-3 (left) and a corresponding increase in soluble LAG-3 upon treatment (right). **(d)** LAG-3 shedding is both dose-dependent (left) and time-dependent (right), with maximal cleavage observed within 6 hours of treatment.

To evaluate induced LAG-3 shedding, we stably transduced HEK293T, HeLa, and Jurkat T cells with FLAG-tagged LAG-3 **(Figure S1)**. None of these parental cell lines exhibited endogenous LAG-3 expression. Upon treatment with the bispecific shedders, we observed robust depletion of cell surface LAG-3 across all cell lines, with LAG-3 shedder 2 demonstrating the highest potency **(Figure 2b)**. Western blot analysis confirmed a dose-dependent reduction of total cellular LAG-3 in both HeLa and Jurkat T cells treated with LAG-3 shedders 1 and 2, while monovalent isotype control arms had no effect. Correspondingly, immunoprecipitation of the cell culture supernatants revealed a concomitant increase in sLAG-3 production following shedder treatment **(Figure 2c, Figure S2)**. This inverse relationship further confirms that target depletion occurs via ADAM10-mediated shedding through induced proximity. Using flow cytometry, we showed that LAG-3 shedding was rapid, with target reduction observed as early as 1 hour and plateaued by 6 hours **(Figure 2d)**.

### Mechanism of LAG-3 induced shedding

To examine the mechanism of LAG-3 depletion, we first visualized the effect of shedder treatment via confocal microscopy. As expected, robust membrane LAG-3 removal by both shedder 1 and shedder 2 were observed, whereas the single-arm isotype controls had no effect **(Figure 3a)**. The disappearance of LAG-3 co-localizes with the plasma membrane, and minimal indication of internalization was observed within the window of treatment. Treatment with Bafilomycin A had no substantial impact on LAG-3 degradation, providing further evidence for a lysosome-independent mechanism of action for the shedders **(Figure S3)**. To quantitatively assay the level of shedder internalization, we labeled the LAG-3 shedders with LysoLight Deep Red (LLDR), a cathepsin-dependent fluorescent probe^34^. LLDR provides a turn-on signal upon cathepsin cleavage in the lysosome **(Figure 3b)**. As a positive control, the fluorescence was compared to a known fast internalizing KineTAC, cetuximab-CXCL12, which efficiently targets EGFR for CXCR7-mediated lysosomal degradation^35^. Consistent with the microscopy images, neither shedder produced substantial fluorescence compared to the KineTAC. Interestingly, LAG-3 shedder 1 showed moderate internalization at high dose with prolonged incubation, suggesting shedding can be combined with internalization to enhance target degradation **(Figure 3c)**. Collectively, these studies confirm that degradation via antibody shedders occurs primarily in the extracellular space and is independent of lysosomal trafficking.

**Figure 3.**
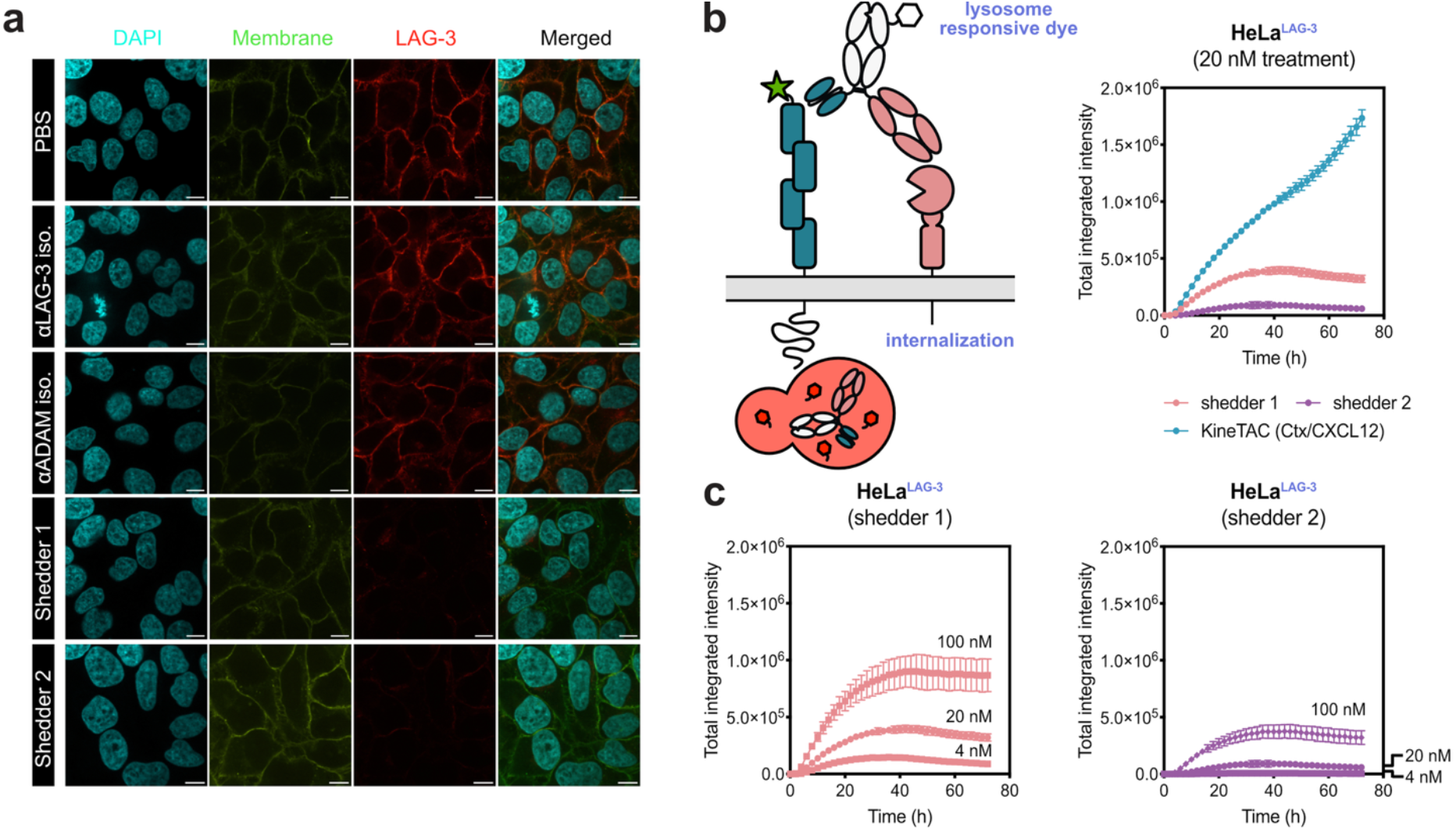
Visualizing induced LAG-3 shedding via microscopy. **(a)** Cells were treated with bispecific shedders or isotype control antibodies for 6 h. Representative immunofluorescence images show robust loss of LAG-3 (red) specifically in shedder-treated cells. LAG-3 signals were localized with the plasma membrane colocalization (yellow) as it disappears during treatment with no clear indication of internalization. **(b)** The loss of LAG-3 is independent of internalization. Cells were treated with bispecific shedders or a known internalizing control (KineTAC) conjugated to a lysosome-sensitive dye. Shedders exhibited minimal internalization signal while the KineTAC control showed robust fluorescence increase over time. Shedder 2 remained in the extracellular space even with prolonged incubation and at high concentrations. **(c)** In contrast, lysosomal entrance was detectable for shedder 1 rather than shedder 2, suggesting that the scFv-Fc format can induce LAG-3 internalization and degradation at high dose. Images were captured every 2 h for 72 h on the Incucyte. Total integrated intensity was calculated by NIRCU x μm2/image. Error bars represented standard deviations for three biological replicates.

Next, we assessed the ADAM10-dependency of induced LAG-3 shedding. ADAM10 expression was partially knocked down in LAG-3-expressing HeLa cells via small-interferring RNA (siRNA). As expected, both depletion of surface LAG-3 and enrichment of sLAG-3 were substantially compromised without ADAM10 **(Figure 4a)**. To confirm the necessity of protease activity, we pre-treated cells with ADAM10 inhibitor GI254023X, which also resulted in a marked reduction in antibody-mediated shedding **(Figure 4b, Figure S4)**. Orthogonally, we generated a LAG-3^ESCP^ mutant containing a 12 amino-acid deletion within the connecting peptide, a modification known to confer resistance to ADAM-mediated shedding^29^. In the LAG-3^ESCP^-expressing HeLa cells, induced shedding was completely impaired **(Figure 4c)**, indicating that the cleavage event is both sequence- and conformation-dependent. Interestingly, LAG-3 shedder 1 retained partial degradation activity under pharmacological inhibition. This observation is consistent with our internalization studies, suggesting that shedder 1 has the ability to mediate some levels of lysosomal degradation. This proteolysis-independent degradation was further corroborated via confocal microscopy **(Figure S5)**. We speculate that this mechanistic difference originates from the distinct bispecific formats: LAG-3 shedder 1 (an scFv-scFv bispecific) may enable transient diabody-like interactions, increasing effective valency and driving receptor clustering and internalization, whereas shedder 2 (an scFv-Fab bispecific) does not. Collectively, these mechanistic studies confirm that our bispecific shedders can achieve target depletion independent of endocytic trafficking, relying instead on enzymatic activity of ADAM10.

**Figure 4.**
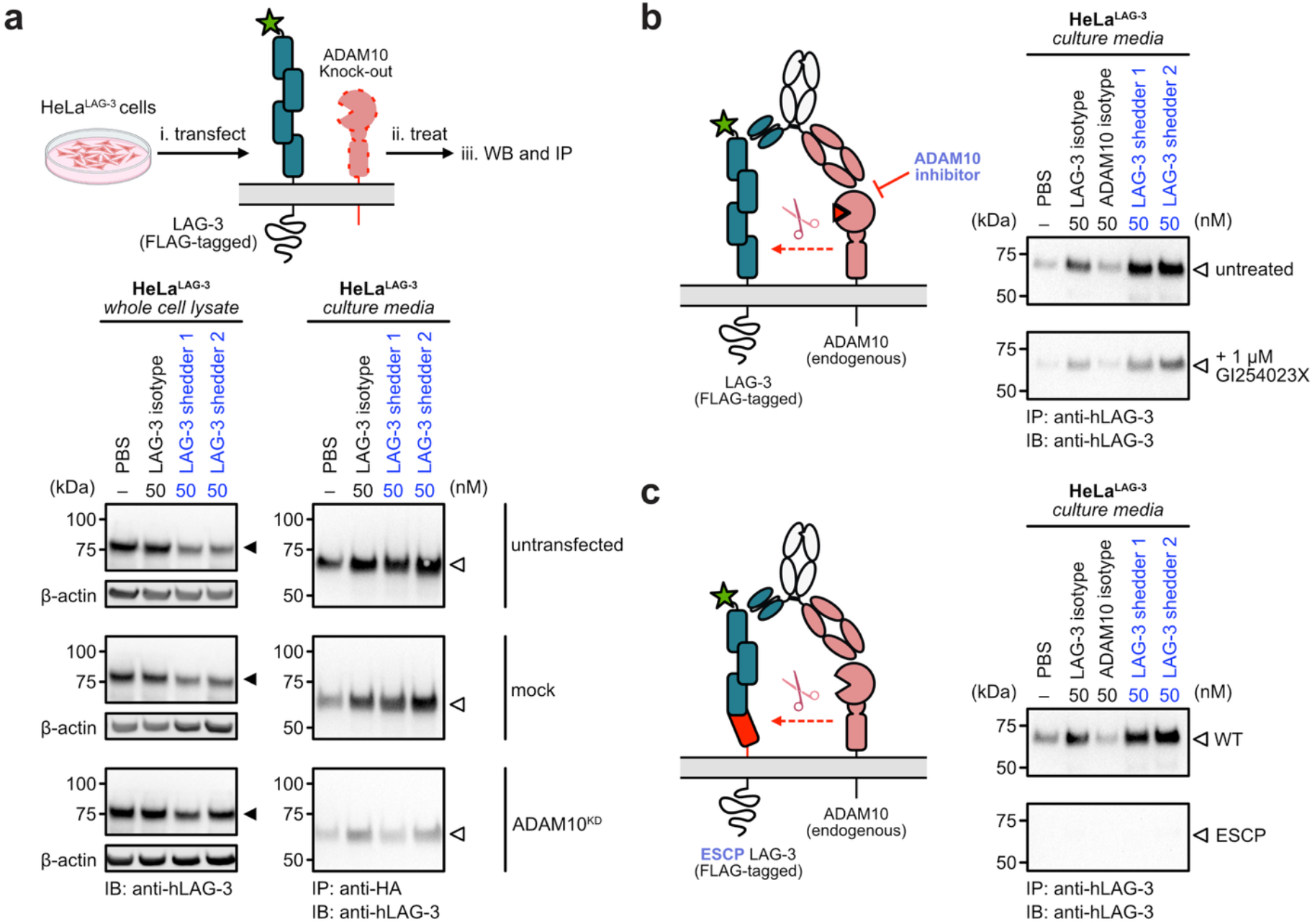
Induced LAG-3 shedding requires ADAM10 proteolysis and intact substrate cleavage site. **(a)** HeLa cells expressing LAG-3 were transfected with ADAM10-targeting or non-targeting (control) siRNA. Western blot analysis shows that ADAM10 knockdown (ADAM10^KD^) prevents the loss of full-length LAG-3 (solid triangle) when treated with the shedders (left). This rescue of cellular LAG-3 also correlated with a significant reduction in the level of soluble LAG-3 (open triangle) in the culture media of ADAM10^KD^ cells. **(b)** Pharmacological inhibition of ADAM10 reduces LAG-3 shedding. LAG-3-expressing HeLa cells were pre-treated with the ADAM10 inhibitor GI254023X (1 µM) for 16 h. Subsequent treatment with shedders resulted in significantly decreased levels of soluble LAG-3 in the culture media compared to vehicle-treated controls, confirming that ADAM10 protease activity is necessary for shedder-mediated LAG-3 release. **(c)** HeLa cells expressing a cleavage-resistant LAG-3 mutant with a 12-amino-acid truncation within the connecting peptide remain non-responsive to shedder treatment. These data further corroborate a mechanism requiring site-specific proteolysis, as the absence of the cleavage motif completely abrogates shedder-mediated LAG-3 depletion.

### LAG-3 shedders potentiate T cell activation

We next evaluated the efficacy of induced LAG-3 shedding in primary human T cells. T cells were isolated from peripheral blood mononuclear cells (PBMCs) and activated with anti-CD3/anti-CD28 antibodies and IL-2 for 48 hours to induce endogenous LAG-3 expression^36^. Following activation, cells were treated with bispecific shedders for 6 hours. Both full-length surface LAG-3 and endogenous shed fragments were detectable in activated T cells, consistent with the known upregulation of ADAM10 and ADAM17 activity during T cell activation. Notably, treatment with Relatlimab resulted in a slight increase in surface LAG-3 expression possibly by stabilizing it at the cell surface. In contrast, both LAG-3 shedders 1 and 2 effectively reduced the total cellular of full-length LAG-3 levels **(Figure 5a)**. This depletion was robust even at concentrations as low as 5 nM, indicating the high potency of the bispecific constructs. Immunoprecipitation analysis of cell-culture supernatants revealed that LAG-3 shedder 2 generated slightly higher levels of sLAG-3 than shedder 1 **(Figure 5b)**, further supporting its superior shedding efficiency, whereas shedder 1 partially functions through receptor internalization.

**Figure 5.**
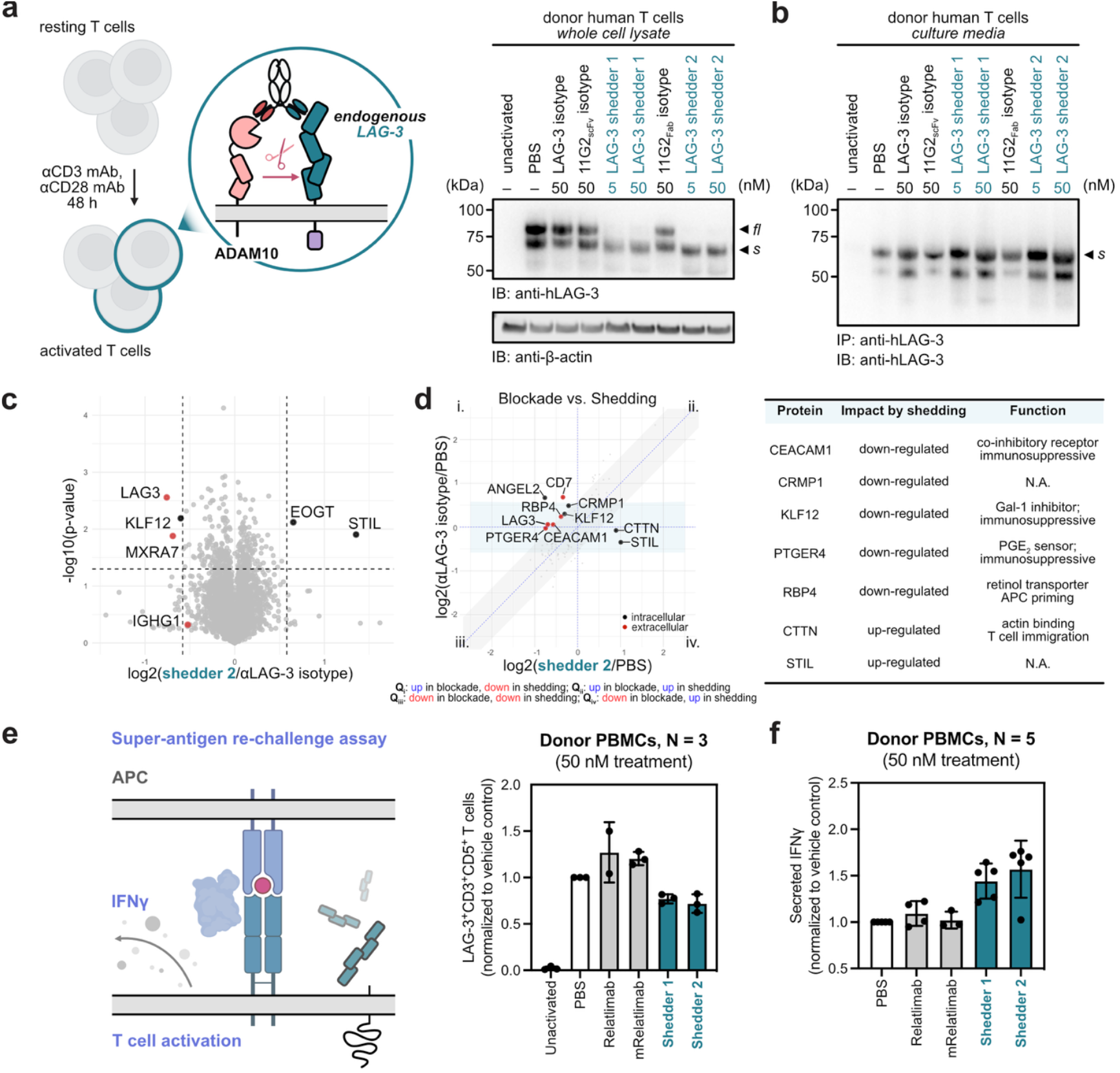
Induced shedding of LAG-3 in primary T cells down-regulates some immunosuppressive proteins and increases IFN*γ* secretion. **(a-b)** Human T cells were activated with a cocktail of antibodies (1 µg/mL anti-CD3/CD28) and IL2 (300U/mL) to upregulate endogenous LAG-3 expression. Cells were then treated with the bispecific shedder or control antibodies. Western blot analysis showed robust depletion of full-length LAG-3 in whole-cell lysates **(a)** and enrichment of soluble LAG-3 in conditioned media **(b). (c-d)** Proteomic profiling distinguishes LAG-3 shedding from antibody blocking. Mass spectrometry analysis of whole-cell lysates from activated T cells reveals a distinct proteomic signature following shedder treatment **(c)**. Shedder treatment induces a significant depletion of LAG-3 while ADAM10 levels remain relatively unchanged. Comparative analysis between cells treated with shedders or a single-arm anti-LAG-3 blocking antibody identifies unique alterations in immune signaling proteins **(d)**. These data suggest that LAG-3 cleavage potentially triggers downstream signaling effects distinct from those of traditional receptor blockade. **(e-f)** LAG-3 shedding enhances activation of primary T cells. LAG-3 expression was induced in donor PBMCs using bacterial superantigen (SEB, 250 ng/mL). Activated cells were treated with LAG-3 shedders, Relatlimab, or control antibodies for 6 h, followed by a second SEB stimulation (5 ng/mL). Flow cytometry analysis confirmed the expected downregulation of surface LAG-3 on CD3^+^CD5^+^ T cells **(e)**. IFN*γ* secretion was quantified via ELISA from the culture supernatants. Shedder-treated cells exhibited significantly enhanced IFN*γ* secretion compared to both isotype controls and Relatlimab-treated cells **(f)**. This elevation in effector function suggests that shedder-mediated LAG-3 depletion may provide a more effective functional enhancement than conventional checkpoint blockade. Error bars represented standard deviations for four biological replicates.

To confirm LAG-3 shedding at the proteomic level, we performed mass spectrometry on whole-cell lysate from activated human T cells following treatment with bispecific shedders, single-arm isotype control, and PBS. Quantitative analysis of the whole-cell lysates revealed that LAG-3 was the most significantly downregulated protein upon shedder treatment, while ADAM10 levels remained unchanged **(Figure 5c)**. To gain insight on potential differences between blockage versus shedding, we compared changes at the proteome level from Relatlimab (relative to PBS) and LAG-3 shedder (relative to PBS) treatments. Protein changes found in both conditions likely represent the consequences of LAG-3 signaling blockade (diagonal zone, **Figure 5d**). Conversely, proteins appearing along the horizonal zone represented changes unique the shedder-treated samples a distinct shedding-centric signature. Intriguingly, several immunosuppressive proteins – including CEACAM1, KLF12, PTGER4, and RBP4 – were uniquely downregulated following induced LAG-3 shedding **(Figure 5d)**. Conversely, we observed the upregulation of CTTN, which control actin-dependent immune cell migration, and STIL. These findings suggest that the proteolysis of LAG-3 may modulate or potentiate T cell function through secondary signaling mechanisms that are fundamentally distinct from traditional checkpoint blockade.

We next assessed the functional consequence of induced LAG-3 shedding in primary human T cells. PBMCs were stimulated with the superantigen Staphylococcal enterotoxin B (SEB), which crosslinks MHC Class II and the T-cell receptor (TCR) to induce polyclonal activation^37^. In the absence of activating agents, no LAG-3 expression was detected regardless of antibody treatment. However, upon activation, we observed robust LAG-3 shedding in T cells and PBMCs following treatment (**Figure 5e, Figure S6**). Interestingly, a second, lower molecular weight cleavage product was prominently observed in the PBMC shedding experiments (**Figure S6**). This suggests that a secondary proteolytic event initiated by ADAM10 might occur in a multi-cellular mixture, which is equally enhanced by antibody-mediated induced proximity. To evaluate T cell effector function^38^ following LAG-3 shedding, we re-stimulated the treated cells with a second dose of SEB and measured IFN*γ* secretion via ELISA. Cells treated with the shedders exhibited significantly enhanced IFN*γ* production compared to both single-arm isotype controls and the Relatlimab **(Figure 5f, Figure S7)**. Collectively, these results corroborate our proteomic data and demonstrate that induced LAG-3 shedding effectively reinvigorates T cell activation.

### Universal induced shedding with anti-ALFA-tag bispecific antibodies

To extend the broad utility and generality of induced shedding, we established a cellular assay to evaluate substrate compatibility. We developed a “universal shedder” based on a high-affinity nanobody that recognizes the ALFA-tag^39^. By using a common epitope tag to recruit diverse POIs, this platform provides a streamlined, turnkey method to examine induced shedding across a variety of targets. To validate this platform, we transfected HeLa cells with either LAG-3 or another canonical ADAM10 substrate, interleukin-6 receptor alpha (IL-6Rα)^40^. Each construct was engineered with an N-terminal ALFA-tag for recruitment and an HA-tag for downstream purification and detection. Western blot analysis confirmed efficient expression of the transgenes and all components required for induced shedding **(Figure 6a)**. Consistent with our results for the anti-LAG-3 shedders, the culture media from cells treated with the ALFA-shedder were significantly enriched with soluble fragments of the targeted transgenes **(Figures 6b, 6c)**. The ALFA-tag induced shedding was also compatible in other cell lines and restricted to protease activity **(Figure S8)**. These results suggest that this transfection-based assay is a versatile and translatable method for evaluating the shedding potential of diverse cell-surface substrates.

**Figure 6.**
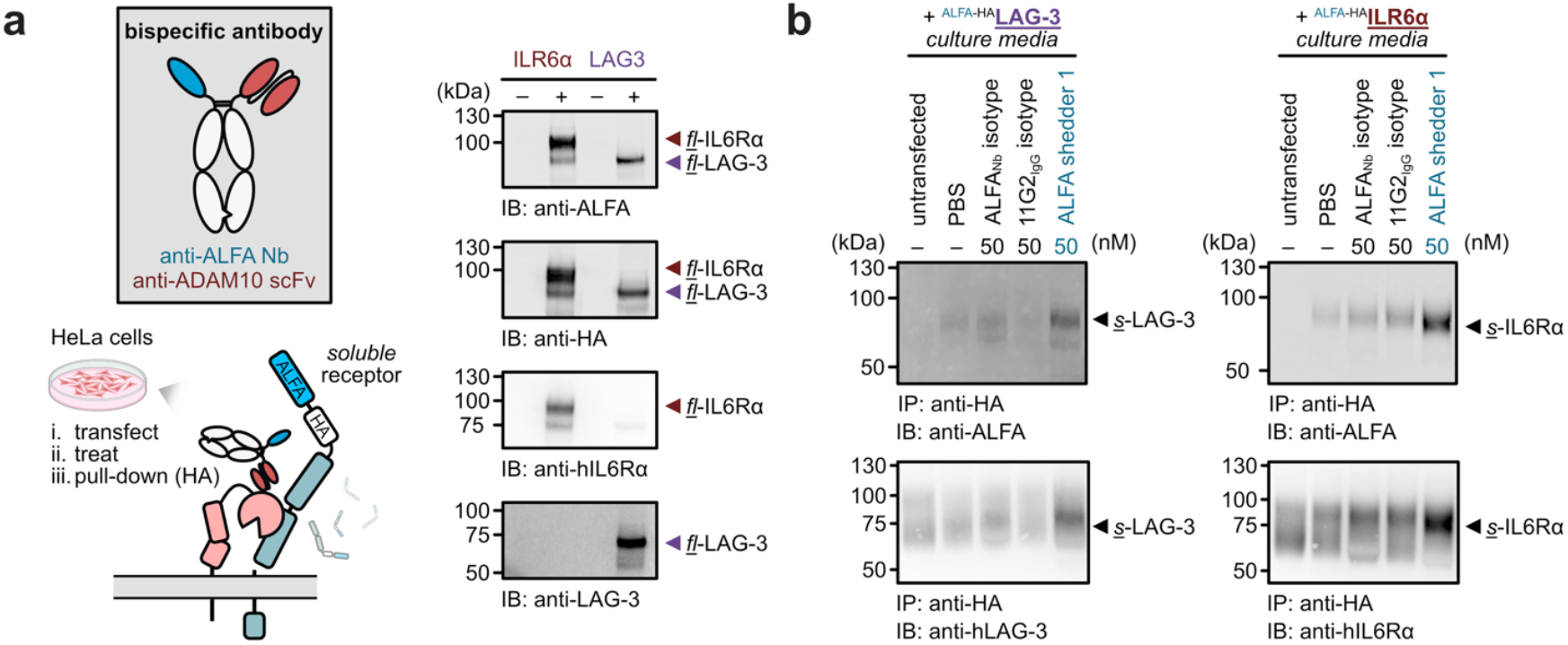
A universal assay for evaluating induced shedders fused with an ALFA tag on the POI substrate. **(a)** Cells were transfected with a POI, either IL6Rα or LAG-3, harboring tandem N-terminal ALFA and HA tags. The ALFA-tagged substrate is targeted by a bispecific shedder utilizing an anti-ALFA nanobody as the antigen-engaging arm. Following treatment, cleaved ectodomain fragments were isolated from the culture media via HA-tag immunoprecipitation (IP) and analyzed to reveal induced shedding of the POI. Western blot analyses confirming proper expression of all transgene components for test substrates LAG-3 and IL6Rα. **(b-c)** ALFA-shedders facilitate proteolysis of LAG-3 **(b)** and IL6Rα **(c)** in the universal shedding assay. Transfected cells were treated with the bispecific shedder or single-arm controls for 6 h. Immunoprecipitation of culture media via the HA-tag reveals the presence of soluble, ALFA-tagged cleavage products (top panel). Immunoblotting confirms these shed fragments as soluble LAG-3 and IL6Rα, respectively (bottom panel). These results demonstrate the modularity of the ALFA-tag system for inducing the shedding of diverse cell-surface substrates.

Leveraging the universal ALFA-shedder, we evaluated whether induced shedding could serve as a platform to investigate signaling events initiated by extracellular proteolysis. It is well-established that the cleavage of IL-6Rα generates a soluble fragment that can form a potent complex with IL-6^41,42^. This “trans-signaling” complex recruits signal-transducer gp130 and triggers downstream signaling cascade on cells that do not express high level of IL-6Rα^42^. We hypothesized that sIL-6Rα generated via induced shedding would remain bioactive and competent in facilitating IL-6 trans-signaling. To test this, we treated cells expressing ALFA-tagged IL-6Rα with the universal shedders and collected the conditioned media. This supernatant was supplemented with recombinant IL-6 and applied to HeLa cells with low endogenous IL-6Rα expression **(Figure S9)**. Gratifyingly, cells treated with supernatant containing the induced shedding products exhibited high levels of STAT3 phosphorylation. These results demonstrate that the cleavage products generated by bispecific shedders are biologically active and capable of recapitulating native signaling roles.

We further utilized the universal shedding assay to probe the epitope preference of our bispecific shedders. For LAG-3, the clinical antibody Relatlimab recognizes an epitope that is spatially proximal to the N-terminal ALFA-tag. Conversely, we developed an anti-IL-6Rα shedder (based on Tocilizumab) that binds to a region distal from the N-terminus^43^. Notably, both the ALFA-tag-mediated and antibody-based shedders exhibited comparable efficacy in soluble fragment release **(Figures 7a, 7b)**. The geometric requirements for POI-engagement are likely flexible, suggesting a broad range of substrates could potentially be targeted for shedding. Next, we assessed the epitope preference of the ADAM-recruiting arm by generating an ALFA-shedder with an ADAM10 binder (8C7) that recognizes a distinct epitope^32^. Consistent with the 11G2-based construct, the 8C7-ALFA shedder competently induced the shedding of IL-6Rα **(Figure 7c)**. Interestingly, 8C7 is known to enhance the catalytic activity of ADAM10 with synthetic peptide substrates. This activation, however, was not recapitulated in this context. Nevertheless, the ability to engage a different ADAM10 epitope while maintaining efficacy further underscores the generality of this platform. Finally, we demonstrated the versatility of the universal shedding system by successfully targeting two additional substrates, MICA^44,45^ and CD62L^46^ **(Figure 7d)**.

**Figure 7.**
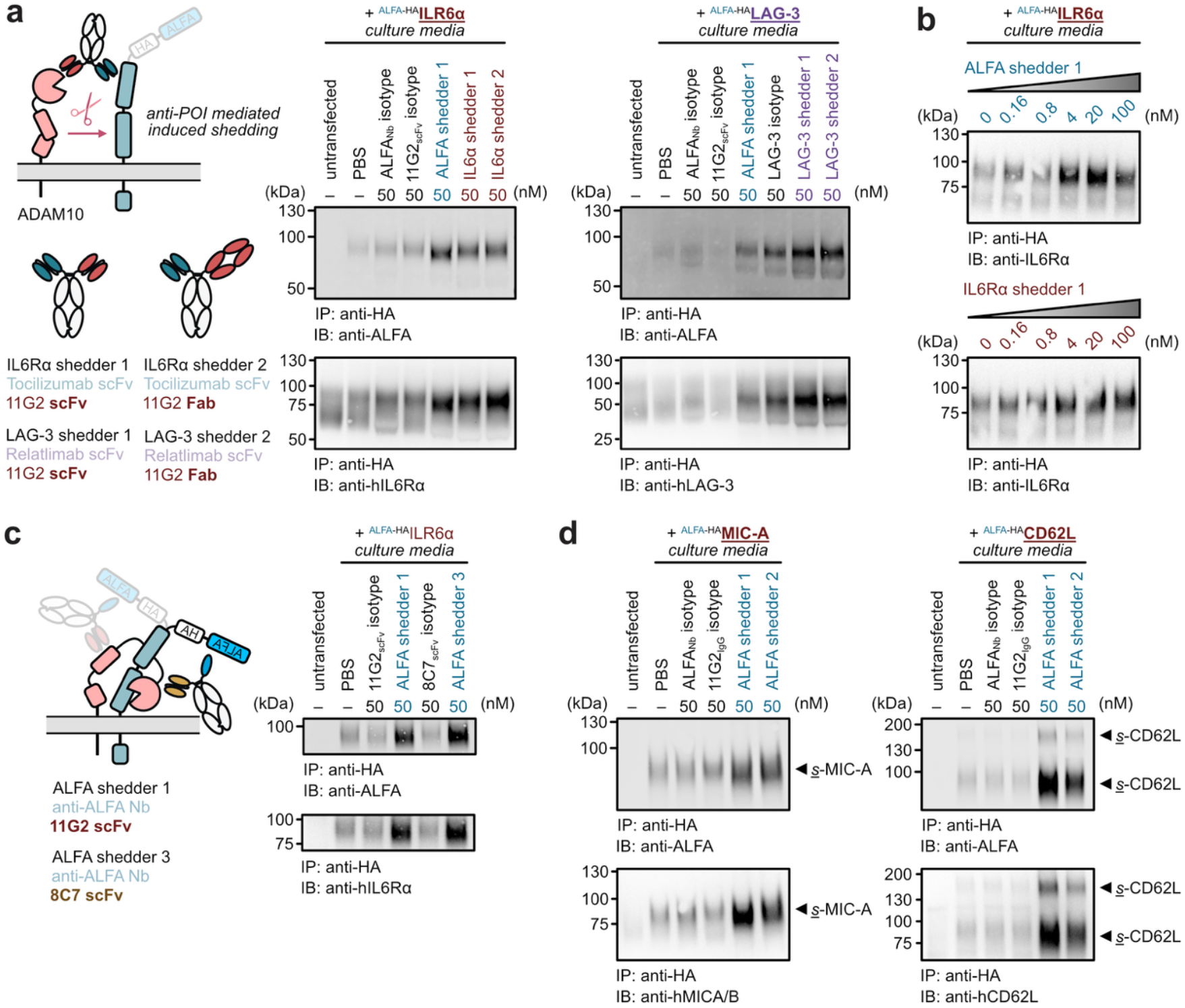
Epitope preference for induced shedding. **(a)** Target engagement via N-terminal binding is sufficient for ADAM10-induced shedding. Diagrams of cells transfected with either ALFA-tagged LAG-3 or IL6Rα and treated with the ALFA-shedder or their respective native antibody-based shedders for 6 h. Following HA-immunoprecipitation and subsequent immunoblotting (right), no substantial differences in soluble fragment production of POIs were observed between the two binding modalities. **(b)** Both ALFA-tag and Tocilizumab-based shedders produce soluble IL6Rα in a dose-dependent manner. Like single-dose analysis, no distinct differences were observed between the two target-binding model suggesting N-terminal targeting is sufficient to facilitate induced shedding. **(c)** Binding to the ADAM10 metalloprotease domain also facilitates induced shedding. A second ALFA-shedder was engineered utilizing the 8C7 clone, which targets the ADAM10 metalloprotease domain proximal to the active site. Comparable levels of soluble fragment production were detected between the 8C7-based shedder and the 11G2-based ALFA shedder. These results demonstrate that target-protease recruitment remains effective across multiple binding epitopes within the ADAM10 extracellular domain. **(d)** The universal shedding assay was extended to additional known ADAM10 substrates. Cells were transfected with ALFA-tagged MIC-A (MHC-I related protein, left) and CD62L (cell adhesion molecule, right). Treatment with the ALFA shedder followed by IP and immunoblotting revealed successful shedding for both targets.

Finally, we demonstrated the broad substrate scope of induced shedding by targeting receptors beyond immunological contexts. We focused on the low-density lipoprotein receptor (LDLR)^47,48^ and LDLR-related protein 8 (LRP8)^49^, both are well-characterized substrates for ADAM10. Proteolytic cleavage regulates the ability of LDLR to bind its cognate ligand, LDL. For LRP8, extracellular cleavage is a prerequisite for the subsequent release of its intracellular domain into the cytoplasm for downstream signaling^50^. We generated bispecific shedders against LDLR and LRP8 respectively, and evaluated their efficacy. Treatment with the respective shedders resulted in a substantial reduction of the target proteins from the cell surface **(Figures S10-11)**. These results demonstrate that our platform enables the targeted extracellular shedding of diverse receptors via induced proximity.

## DISCUSSIONS

Extracellular targeted protein degradation (eTPD) has emerged as a powerful modality for the selective removal of cell-surface proteins^2^. Plethora of modalities now exist to recruit different recycling receptors or membrane-tethered E3 ligases to traffic POI from outside of the cell to inside for degradation^5,6,8,10,35^. In contrast to lysosomal proteolysis which leads to complete protein turn-over, extracellular proteolysis is generally targeted and site-specific. Proteomic studies have revealed a plethora of site-specific proteolytic events on the cell surface leading to nicked and shed proteins but most without functional annotation. Here, we provide a tool for target specific proteolysis and shedding for cell surface proteins. Induced shedding offers a new paradigm for recruiting membrane-resident proteases to cleave a POI directly at the cell surface. Mechanistically, the shedders are faster than lysosomal trafficking-based degraders, as they leverage a direct enzymatic reaction rather than complex multi-step endocytic pathways to deliver to the lysosome and subsequent proteolysis. We show that *Shedders* recruiting ADAM10 enables efficient, selective, and proteolysis-dependent shedding of the immune checkpoint receptor LAG-3, as well as other therapeutically relevant membrane proteins. In primary T cells, LAG-3 shedders outperform analogous degraders based on recycling receptors CXCR7 or IL7R^35^, highlighting the rapid nature of proteolysis-based mechanism **(Figure S12)**.

The initial *Shedder* design was guided by two core principles: (1) a high-affinity POI-engaging arm and (2) an innocuous ADAM10-recruiting arm. High-affinity POI binding ensures that shedding occurs with high specificity upon protease recruitment. Notably, our LAG-3 shedders demonstrated high potency at low concentrations, although it’s unclear whether this induced proteolysis is catalytic. In principle, even greater shedding efficiency could be achieved by “detuning” the affinity of the POI arm. For protease recruitment, we show that a non-perturbing binder is sufficient for recruiting a constitutive sheddase like ADAM10. However, many other cell-surface proteases exist in autoinhibited states^52–54^. Consequently, the use of conformation-specific or allosteric modulatory binders represents a promising strategy to further enhance shedding potency when recruiting proteases with more complex regulatory mechanisms.

A key feature of induced shedding is its ability to release bioactive fragments into the extracellular milieu. For therapeutic targets such as LAG-3, this provides an additional layer to elicit pharmacological advantage over traditional degradation or antibody-mediated blockade^55^. Specifically, we demonstrated that LAG-3 shedders potentiate T cell function through mechanisms distinct from the simple receptor-blockage provided by Relatlimab. This was further reflected in a superantigen rechallenge assay, where shedders outperformed Relatlimab in rejuvenating the ability of activated T cells to secrete IFN*γ*^38^. While the exact mechanism underlying this enhanced effect remains to be fully elucidated, our proteomic analysis revealed unique alterations in regulatory proteins critical for immune activation. Future work will further characterize the role of LAG-3 shedding in modulating T cell activation and explore its application in other immunotherapy contexts, such as CAR-T cell therapy^56^. Similar to donor T cells, we observed that LAG-3 expression induced on CD19 CAR-T^57^ cells upon target cell engagement could be similarly depleted via the bispecific shedders **(Figure S13)**. In the case of IL-6Rα, we also showed that the shed extracellular domain remains competent to form active signaling complexes with soluble cytokines. This dual capability of simultaneous surface target depletion and fragment release opens the possibility of utilizing induced shedding as a platform for on-demand delivery of therapeutically relevant molecules at the cell surface.

Finally, we anticipate that shedders will serve as valuable tools for studying specific extracellular proteolysis events. Investigating discrete cleavage events at the cell surface remains a significant challenge due to the inherent promiscuity of membrane proteases. Furthermore, neither pharmacological inhibition nor genetic knockdown is sufficient to fully decouple the specific functional consequence of proteolysis from broader cellular changes. Induced shedding thus provides a targeted, gain-of-function approach to selectively upregulate individual proteolytic event. This strategy not only recapitulates the protease-mediated removal of a target protein but also preserves the biological signaling potential of the shed soluble extracellular domain and the membrane-tethered stub. In summary, the induced shedding modality represents a transformative paradigm for hijacking cell-surface proteases, offering a versatile toolkit for both mechanistic investigation and next-generation therapeutic protein degradation.

## ACKNOWLEDGMENT

We thank Dr. Kaitlin Schaefer, Dr. Brandon Holmes, and Dr. Jun Wang for their assistance and helpful discussions as well as the entire Wells lab for helpful discussion and expertise. We thank Dr. Arun Witta, Dr. Alex Marson, and members of the Witta and Marson labs for insights in CAR-T cells transduction and CD19-targeting CAR plasmid. We are grateful to generous support from NIH-1R01CA248323-01(J.A.W), NIH-R35GM122451 (J.A.W.), and the Hind Professorship in Pharmaceutical Sciences (J.A.W). Z.Y. and T.M.P.C are supported by a National Institute of General Medical Sciences F32 Postdoctoral Fellowship. F. Z. is supported by a postdoctoral fellowship funded by the A.P. Giannini Foundation. K. K. is supported by a graduate fellowship funded by the National Science Foundation.

## AUTHOR CONTIRBUTIONS

Z.Y., F.Z., and J.A.W conceived and designed the study. Z.Y. and F.Z. designed and characterized bispecific shedders. Z.Y., F.Z., Y.Z., S.G., Y.W. and K.K. cloned and expressed the recombinant proteins. Z.Y. and F.Z. performed the western blot and immunoprecipitation experiments in engineered and primary cell lines. A.M. performed the siRNA knock-down on LAG-3 expressing HeLa cells K.M. performed the microscopy experiments. T.M.P.C. performed the proteomic experiments. Z.Y. and F.Z. performed the functional characterization of shedding. Z.Y., F.Z., and K.M. developed the universal shedding assays and characterized additional substrates for induced shedding. Z.Y., F. Z., and J.AW. wrote the manuscript and all authors reviewed and edited the manuscript.

