## Supplemental File for "Targeted shedding of extracellular membrane proteins by induced protease recruitment"

#### MATERIALS AND METHODS

*Plasmid construction*

All the IgGs were constructed in a pcDNA3.4 vector that expresses the light chain and heavy chain, respectively for mammalian expression. For generating bispecific antibody, the heavy chain variable regions or scFvs were cloned into zymework-A mutant Fc and zymework-B Fc sequences respectively. The anti-ADAM10 11G2 and anti-LAG3 Relatlimab antibody variable regions were cloned into human IgG1 expression construct with LALAPG mutants to abolish Fc effector function.

*Protein expression*

Antibodies were expressed in Expi293F cells on a 30 mL scale. In brief, 24 µg of DNA was added to 3 mL of OptiMEM, followed by 24 µL of the FectoPro transfection reagents. After 10 min of incubation, 27 mL of Expi293F cells at 3 million/mL was added and shaken at 37 °C. On the second day, 300 µL of 300 mM vaporic acid and 270 µL of 45% glucose were added to the cells. After 5 days of transfection, cells were harvested, spun down at 4000g for 20 min, and filtered by 0.45 µm steri-flip. Supernatants were then incubated with Sepharose A resin for 2 h, and proteins were then eluted by 0.1 M acetic acid and neutralized by Tris pH 11. Proteins were buffer changed 3 times in PBS in Amicon tubes. Purity and integrity of all proteins were assessed by SDS–PAGE.

*Cell lines*

HeLa (ATCC), HEK293T (ATCC), MDA-MB-231 (ATCC) cells were cultured in and Lenti X-293T cells (Takara no. 632180) were cultured in complete medium: DMEM (ThermoFisher Scientific) containing 4.5 g/L D-glucose, 2 mM L-glutamine, and supplemented with 10% (v/v) FBS (Life Technologies), penicillin (100 U mL−1) and streptomycin (100 µg mL−1, Gibco). Jurkat (ATCC), Raji (ATCC), SK-ND-Z (ATCC), and THP-1 (ATCC) cells were cultured in RPMI-1640 (ThermoFisher Scientific) with 10% (v/v) FBS and penicillin (100 U mL−1) and streptomycin (100 µg mL−1). Expi293 cells were cultured in FreeStyle 293F medium (ThermoFisher Scientific, catalog no. A1435103).

*Lentiviral cell line construction*

Stable cell lines were generated by lentiviral transduction. HEK293T Lenti-X cells were transfected with second-generation lentiviral packaging plasmids at approximately 80% confluence. FuGene HD (Promega) was used for transfection. After 72 hours, supernatant was harvested and filtered. Cleared supernatant was added to target cells with polybrene and centrifuged at 1000g at 33°C for 2 hours. Cells were incubated with viral supernatant overnight before the media was changed to fresh complete DMEM. Cells were expanded for 48 hours before being grown in drug-selection media. After 72 hours, cells were analyzed by flow cytometry for expression.

*PBMC and T cell Isolation*

Human peripheral blood mononuclear cells (PBMCs) were isolated from leukoreduction chamber residuals following Vitalant Blood Donation using established protocols. Briefly, PBMCs were isolated using Ficoll separation in SepMate tubes (STEMCELL Technologies) in accordance with the manufacturer's instructions. Total T cells were isolated from PBMCs using the EasySep™ Human T Cell Isolation Kit (STEMCELL), following the manufacturer's protocol. PBMCs and T-cells were maintained in RPMI-1640 with 10% FBS, 1× MEM NEAA, 1 mM sodium pyruvate, 10 mM HEPES, 50 µM 2-mercaptoethanol (sRPMI). Purity was assessed via flow cytometry.

*LAG-3 shedding assays with engineered cell lines*

Cells stably expressing N-terminal FLAG-tagged LAG-3 were plated in 12- and 24-well plates and grown to 70% confluency before treatment. Jurkat-LAG-3 cells were used at ($0.1 \times 10^6$ per well) in a 12-well plate. Medium was aspirated, and cells were treated with bispecific or control antibodies in complete growth medium. After incubation at 37 °C for 6 hours (or indicated amount of time), the conditioned media was removed and flash-frozen for immunoprecipitation. The cells were lifted with PBS and collected by centrifugation at 500 x g for 5 min at 4 °C. Samples were then lysed for western blotting or analyzed via flow cytometry to quantify protein level.

*LAG-3 shedding assays with donor PBMCs and T cells*

Human PBMCs and T cells were isolated and maintained as previously stated. Cells ($2 \times 10^6$ per well) were plated in a 6-well plate coated with anti-CD3/CD28 (1 μg/mL). For T cell assays, the culture media was supplemented with IL2 (300 U/mL). After 2 days, shedders or control antibodies were added to induced LAG-3 shedding and incubated for another 6 h. At the end of the treatment, culture media were harvested for immunoprecipitation and cells were collected for western blot or flow cytometry analysis.

*Shedding assays with immobilized cell lines*

Cells were plated in 12- and 24-well plates and grown to 70% confluence before treatment. Medium was aspirated, and cells were treated with bispecific or control antibodies in complete growth medium. After incubation at 37 °C for 6 hours (or indicated amount of time), the conditioned media was removed and flash-frozen for immunoprecipitation. The cells were lifted with PBS and collected by centrifugation at 500 x g for 5 min at 4 °C. Samples were then lysed for western blotting or analyzed via flow cytometry to quantify protein level.

*Universal shedding assays with HeLa or HEK293T cell lines*

HeLa cells ($0.1 \times 10^6$) were seeded in 24-well dishes (Corning). Transfections were performed with ALFA-tagged receptor constructs using Lipofectamine 3000 (Thermo Fisher) according to the manufacturer’s instructions. Transfections were performed when cells were 75–80% confluent (1 d post seeding). Approximately 16 h post transfection, cell media were aspirated and replenished with fresh media containing the indicated shedders or control antibodies. Cells were further incubated for 6 h at 37 °C (unless specified), lifted with versene (PBS, $Ca^{2+}$ and $Mg^{2+}$ free with 0.04% EDTA, UCSF cell culture facility), and harvested for Western blot analysis. Culture media were aliquoted and flash-frozen for immunoprecipitation (IP). Prior to IP, Pierce Protein A/G magnetic agarose beads (ThermoFisher Scientific, catalog no 78609) were washed with TBST (1 mL x 3) and pre-coated with anti-HA antibody (1 μg of antibody per 10 μL of beads per sample). Antibody-coated beads were added to culture media and the suspension were further incubated at 4 °C for 16 h. Culture media were removed using a DynaMag-2 magnetic rack (Invitrogen) and the beads were washed with TBST (1 mL x 3) and water.

*RNAi Knockdown of ADAM10*

siRNAs (Dharmacon) of ADAM10 or scramble sequence were transfected to LAG-3 expressing HeLa cells following previously established methods. In short, $2 \times 10^5$ HeLa cells were seeded in a 6-well dish with complete DMEM. The next day, cells were transfected with the target siRNA at a final concentration of 100 nM, using Lipofectamine RNAi/MAX (following manufacturer protocol) in OptiMEM and incubated for 16 h at 37°C with 5% $CO_2$. Following transfection, medium was aspirated, and cells were treated with bispecific or control antibodies in complete growth medium. After incubation at 37 °C for 6 hours (or indicated amount of time), the conditioned media was removed and flash-frozen for immunoprecipitation. The cells were lifted with PBS and collected by centrifugation at 500 x g for 5 min at 4 °C. Samples were then lysed for western blotting or analyzed via flow cytometry to quantify protein level.

*Super-antigen rechallenge assay with PBMCs*

Frozen PBMCs were thawed and allowed to recover in sRPMI at 37 °C overnight. Cells (0.2 x $10^6$) were plated in a 96-well plate and activated with SEB (250 ng/mL). After 48 h, cells were washed with sRPMI (2 x 200 µL) and resuspended in media containing SEB (5 ng/mL), bispecific shedders, and control antibodies. Plates were incubated at 37 °for another 48. At the end of treatment, cells were harvested for flow cytometry analysis. The cultured media were also harvested and flash frozen for secreted IFNγ quantification.

*LAG-3 Shedding assays with CAR T cells after repeated stimulation*

CD19 CAR lentivirus was generated as previously described. T cells were isolated from frozen PBMC using EasySep Human T Cell Isolation kit (Stemcell Technologies) and sub-cultured with 30 U/mL IL2 in sRPMI. Cells were activated with Dynabeads human T-activator CD3/CD28 in sRPMI supplemented with IL7 and IL15. After 2 d, activated T cells were transduced with CD19 CAR lentivirus and maintained in sRPMI with IL7 and IL15 for another 7 d. Expression of CD19 CAR were confirmed via flow cytometry.

CD19 CAR T cells (5 x $10^6$) were collected by centrifugation at 500 x g for 5 min and diluted in sRPMI with IL7 and IL15 (5 mL). Raji cells (5 x $10^6$ in 5 mL, 1:1 E:T) were added, and the mixture was incubated for 2 days. This process was repeated for 4-cycle. At the end of the stimulation, CD19 CAR T cells were isolated via FACS sorting and recovered in sRPMI overnight. The sorted CAR T cells were plated (0.1 x $10^6$ per well) in a 96 well and treated with LAG-3 shedders or control antibodies (10 nM or 100 nM). Raji cells (0.1 x $10^6$ per well) were added to each well, and the plate was incubated for another 48 h. At the end of the treatment, cells were washed with PBS (3 x 100 µL) and analyzed via flow cytometry.

*IFNγ ELISA*

Flash frozen supernatant was thawed on ice and diluted 1:100 or 1:250 with PBS. The amount of secreted IFNγ was quantified via ELISA MAX™ Deluxe Set (Biolegend). In short, 384-well Maxisorp plates were coated with provided Human IFN-γ ELISA MAX™ Capture Antibody overnight at 4 °C and subsequently blocked with BSA (2% w/v) for 1 h at RT. 50uL cell supernatant from above were captured onto pre-coated wells for 1h. After three times of wash, Human IFN-γ ELISA MAX™ Detection Antibody were added to the plates and incubated for 1 h at RT. After three times of wash, 1:1000 diluted Avidin-HRP were added to the plates and incubated for 30 min. After three times of wash, antibody binding was detected by TMB substrate (VWR), quenched by 1 M phosphoric acid, and read at 450 nm. Measured absorptions were quantified using standard curve from kit-provided interferon gamma standard.

*Flow cytometry*

Cells were collected by centrifugation at 500 x g for 5 min. Pellets were washed once with PBS + 1% BSA. Cells were incubated with fluorophore-conjugated antibodies in PBS + 1% BSA for 15 min at RT or 30 min at 4 °C. Cells were washed three times and resuspended in cold PBS for flow analysis. Antibodies used included PE anti-human CD223 (LAG-3) (Biolegend, Cat#369306, 1:400), APC anti-human CD223 (LAG-3) (Biolegend, Cat#369212,1:400), Alexa fluor 647 goat anti-human IgG (H+L) (Invitrogen, Cat#A-21445, 1:1000), Alexa fluor 647-conjugated protein A (Invitrogen, Cat# P21462, 1:1000), M1-FITC (gift from Manglik lab, UCSF), PerCP/Cy5.5 anti-human CD3 (Biolegend, Cat#300429, 1:500), PerCP/Cy5.5 anti-human CD5 (Biolegend, Cat#300619, 1:500), BV421 anti-human CD69 (Biolegend, Cat#310929, 1:500), FITC anti-human CD4

(Biolegend, Cat#344604, 1:500). Dead cell staining included propidium Iodide (Biolegend, Cat#421301, 1:250), and LIVE/DEAD™ fixable violet dead cell stain kit (Invitrogen, Cat# L34964). Flow cytometry was performed using a CytoFLEX cytometer (Beckman Coulter, v.2.3.1.22) and CytoExpert software (v.2.3.1.22). Data were analyzed with FlowJo (v.10.8.0).

*Western blotting*

Cells were lifted with PBS + 0.05% EDTA, transferred to Eppendorf tubes, spun down at 500 g for 4 min, and washed twice with PBS. Cells were lysed with 1× RIPA lysis buffer (EMD Millipore) with cOmplete mini protease inhibitor cocktail (Sigma-Aldrich) at 4 °C for 20 min. Lysates were centrifuged at 20,000g for 10 min at 4 °C to remove debris. Soluble protein concentrations were quantified by Rapid Gold BCA Protein Assay Kit (Pierce). Lysates were mixed with 4× Nupage LDS Sample Buffer (Invitrogen) and 2-mercaptoethanol, and then run on NuPAGE 4–12% Bis Tris Protein Gels (Thermo Fisher Scientific). Proteins were transferred to polyvinylidene difluoride membranes using the iBlot2 Western Blotting Transfer System (Thermo Scientific). Membranes were blocked with TBS + 5% BSA + 0.5% Tween for 1 h, and stained with primary antibodies overnight. After three washes, membranes were stained with secondary antibodies for 1 h at RT.

After three washes, membranes were imaged with a LICOR imager or the ChemiDoc MP imaging system (BioRad). Antibodies used included rabbit anti-human LAG-3 (Cell Signaling Technology, catalog no. 15372S, 1:1000), rabbit anti-human IL6R (Cell Signaling Technology, catalog no. 18935S, 1:1000), mouse anti-human □-actin(Cell Signaling Technology, catalog no. 3700S, 1:2000), goat anti-human LDLR (R&D Systems, catalog no. AF2148, 1:1000), rabbit anti-human CD62L (Cell Signaling Technology, catalog no. 58225T, 1:1000), rabbit anti-human MICA/B (Cell Signaling Technology, catalog no. 64899S, 1:1000), FluoTagX2 anti-ALFA LI-COR IRDye 800CW (NanoTag biotechnologies, catalog no. N1502-Li800-L), IRDye 800CW goat anti-rabbit IgG (LI-COR Biosciences, catalog no. 926-32211), IRDye 680RD goat anti-mouse IgG (LI-COR Biosciences, catalog no. 926-68070, 1:5000), IRDye 800CW donkey anti-goat IgG (LI-COR Biosciences, catalog no. 926-32214, 1:5000), and peroxidase goat anti-rabbit IgG (H+L) (Jackson ImmunoResearch, catalog no. 111-035-144, 1:5000).

*Immunoprecipitation*

For each well of cells, 10 μL Protein A/G agarose magnetic bead (Thermo Fisher Scientific, 78609) slurry was washed 2 times with PBST and incubated with capture antibodies: anti-LAG3 (1 µg per sample, Biolegend, Cat#369302), anti-HA (1 µL per sample, cell signaling technology, Cat#3724T) 30 min at RT or 1 h at 4 °C. Antibody-coated beads were added to supernatant and the suspension were further incubated at 4 °C for 16 h. Culture media were removed using a DynaMag-2 magnetic rack (invitorgen) and the beads were washed with TBST (1 mL x 3) and water (1 mL x 1). Proteins were eluted with glycine buffer (pH = 2) and neutralized with tris buffer (1M, pH = 11).

*Confocal microscopy*

HeLa cells were plated on the chambered coverslip (Ibidi, 8-well uncoated) and incubated for 24 h at 37°C. Cells were then treated with 50 nM bispecific shedder or control antibodies in complete growth medium. After 6 h of incubation at 37°C, cells were stained with Membrite (Biotium, 30094-T) and washed with PBS. Cells on the coverslips were fixed with paraformaldehyde (PFA) for 15 min at RT. After washed 3 times by PBS, the resulting sample were stained with anti-LAG3 rabbit antibody (Biolegend, Cat# 369302) and DAPI (Cell Signaling Technologies). Goat anti-mouse IgG 647 (Invitrogen, Cat#A-21240) were stained for visualization. Samples were imaged using a Nikon Ti Microscope with a Yokogawa CSU-22 spinning disk confocal and a 60x objective lens; 405-, 488- and 647-nm lasers were used to image DAPI, LAG-3 and Membrite, respectively. Images were deconvoluted and processed using NIS-Element (v5.21.03) and Fiji software (v2.1.0) packages.

*Lysolight Deep Red Assay*

Antibodies were labeled using the LysoLight Antibody Labeling Kits (Invitrogen, cat. no. L36003) following manufacturer's instructions. Briefly, antibodies were labeled with LLDR with a molar ratio of 1:6 in the presence of 100 mM sodium bicarbonate (pH 8.4) for 2 h at RT. Antibodies were then purified with 7k Zeba dye and biotin removal columns (Thermo Scientific, catalog no. A44297). Cells were seeded at 5000/well on a 96-well polystyrene tissue culture treated plate (Corning, catalog no. 3596). The next day, media was removed, treated with LLDR-labeled antibodies, and then imaged on the Incucyte every 2 h for 72 h.

Degradation Experiments

Cells were plated in 6- or 12-well plates and grown to ~70% confluency before treatment. On the next day, cell culture medium was aspirated, and various concentrations of antibodies in 1 mL of culture medium were then added to each well. Cells were incubated for 24 h at 37 °C prior to flow cytometry or Western blotting experiments.

*Mass Spectrometry*

For proteomic analysis, T cells were isolated from frozen PBMC using EasySep Human T Cell Isolation kit (Stemcell Technologies) and sub-cultured with 30 U/mL IL2 in sRPMI. Cells were activated (2 x 106 per well) with 300 U/mL IL2 on a 6-well plate coated with anti-CD3/CD28 (1 µg/mL) for 48 h, followed by treatment with bispecific shedders or control antibodies (50 nM) for 6 h. Approximately 2 million treated T cells were washed with PBS (3 x 1 mL). Cell pellets were lysed with a 2× dilution of commercial RIPA buffer (Millipore) supplemented with 1× protease inhibitor cocktail (Sigma) and 2 mM EDTA (Sigma) for 10 min at 4 °C. Cells were further disrupted with probe sonication (20% amplitude, 5 min, 4 °C), followed by cell debris removal (20,000g, 10 min, 4 °C), and the clarified cell lysates were then incubated with 50 µL of high-capacity NeutrAvidin-coated agarose beads (Thermo) in Poly-Prep chromatography columns (Bio-Rad) for 2 h at 4 °C to isolate biotinylated glycoproteins. To enrich for biotinylated proteins, the resin was washed sequentially with 5 mL of 1× RIPA (Millipore) plus 1 mM EDTA, 5 mL of high-salt PBS (20 mM phosphate (pH 7.4) with 1 M NaCl (Sigma)) and 5 mL of denaturing urea buffer (50 mM ammonium bicarbonate and 2 M urea). Proteins on the beads were next reduced, carbidomethylated, digested, and desalted using the Preomics iST mass spectrometry sample preparation kit (Preomics) per the manufacturer's recommendations. After desalting, samples were dried, resuspended in 0.1% formic acid, and quantified using the Pierce peptide quantification kit (Thermo Scientific) before liquid chromatography–tandem mass spectrometry analysis.

Liquid chromatography–tandem mass spectrometry was performed by using a Bruker NanoElute chromatography system coupled to a Bruker timsTOF Pro mass spectrometer. Peptides were separated using a prepacked IonOpticks Aurora (25 cm × 75 µm) C18 reversed-phase column (1.6-µm pore size, Thermo) fitted with a CaptiveSpray emitter for the timsTOF Pro CaptiveSpray source. For all samples, 200 ng of resuspended peptides was injected and separated using a linear gradient of 2–23% solvent B (solvent A: 0.1% formic acid and 2% acetonitrile; solvent B: acetonitrile with 0.1% formic acid) over 90 min at 400 µL min 1 with a final ramp to 34% B over 10 min. Separations were performed at a column temperature of 50 °C. Data-dependent acquisition was performed using a timsTOF PASEF tandem mass spectrometry method (TIMS mobility scan range of 0.70–1.50 V•s cm2, mass scan range of 100–1700 m/z, ramp time of 100 ms, 10 PASEF scans per 1.17 s, active exclusion of 24 s, charge range of 0–5 and minimum MS1 intensity of 500). The normalized collision energy was set at 20.

### EXTENDED DATA AND FIGURES

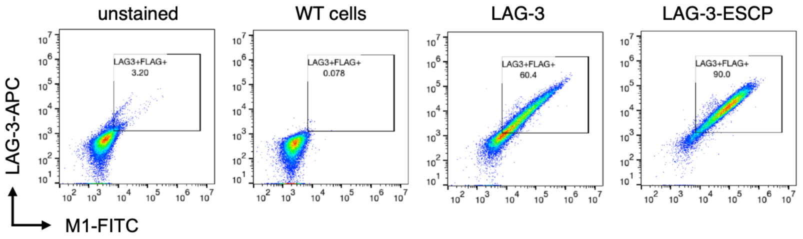


**Figure S1. Engineering LAG-3 expression in model cell lines.** HEK293T cells were transduced with full length LAG-3 bearing N-terminal FLAG-tag via lentivirus. Surface expression of LAG-3 was confirmed via flow cytometry staining for both LAG-3 and FLAG-tag. A control cell line with a cleavage resistant LAG-3 variant was generated by deleting the 12 amino acids connecting peptide on the membrane proximal region.

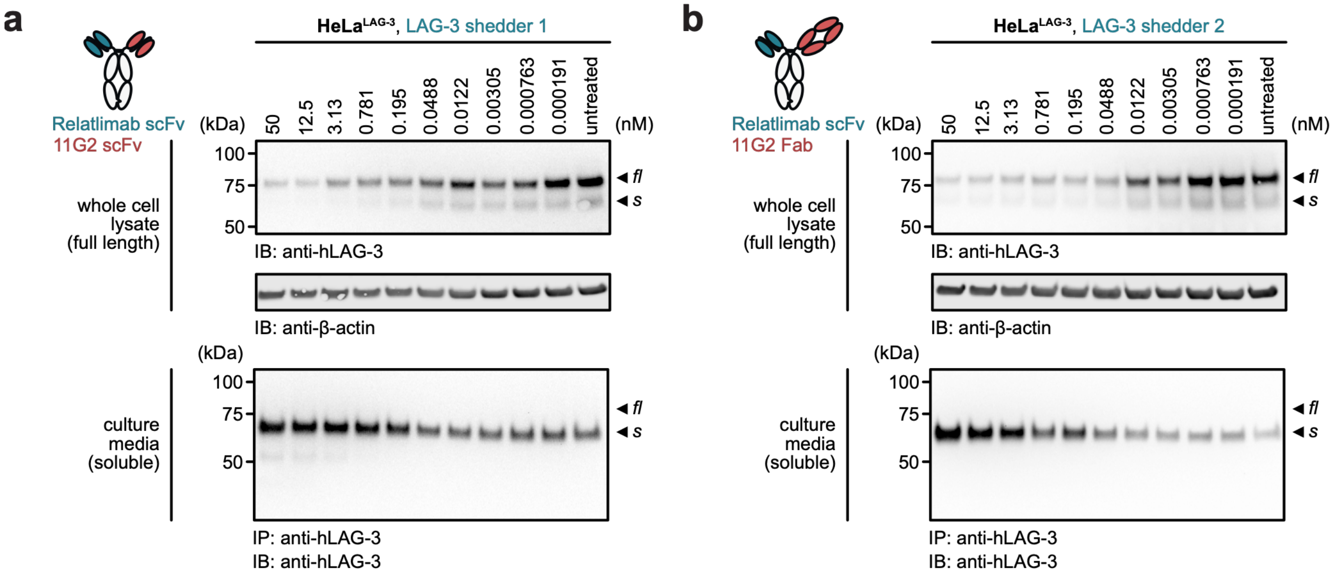


**Figure S2. Induced LAG-3 shedding in engineered HeLa cells.** Cells were treated with increasing concentrations of either LAG-3 shedder 1 **(a)** or shedder 2 **(b)** for 6 h. (Top panel). Western blot analysis of the whole cell lysate revealed the degree of full-length LAG-3 degradation (top panel), while immunoprecipitation of the culture media reveals the amount of soluble LAG-3 production (bottom panel). For both shedders, dose dependent degradation was observed with sub-nM potency. The loss of full-length LAG-3 was reciprocally mirrored in soluble LAG-3 production confirming a shedding-based mechanism.

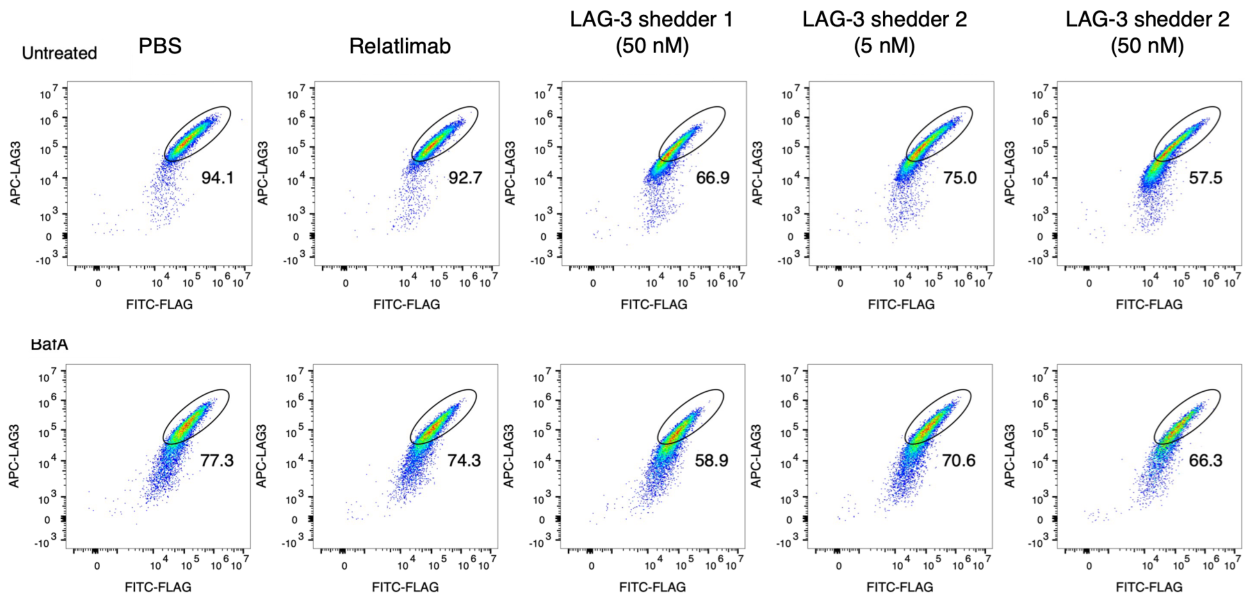


**Figure S3. Induced LAG-3 shedding is independent of lysosomal trafficking.** HeLa cells expressing LAG-3 were pre-treated with Bafilomycin A1 (BafA, 50 nM) for 1 h, followed by treatment with LAG-3 shedders for 6 h. Flow cytometry analysis revealed BafA treatment slightly altered baseline surface LAG-3 level, likely due to interference with endogenous receptor trafficking. However, the loss of LAG-3 mediated by induced shedding was insensitive to BafA. The cell population was gated sequentially on cells, singlets, and live events, and then on LAG-3–APC and FLAG-FITC double-positive cells.

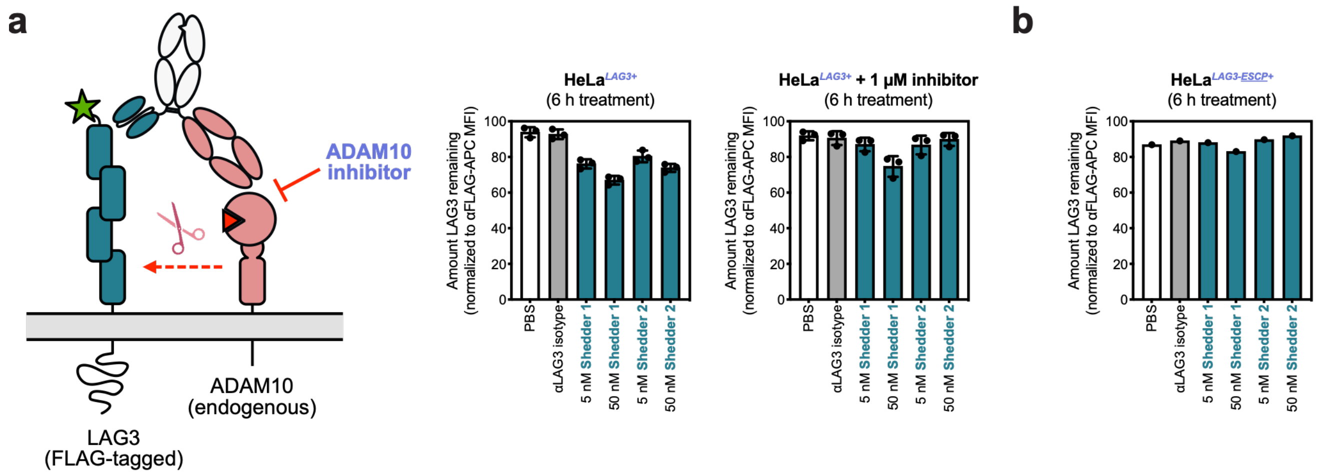


**Figure S4. Induced LAG-3 shedding necessitates ADAM10 proteolytic activity. (a)** HeLa cells expressing LAG-3 were pre-treated with ADAM10 inhibitor GI254023X (1 μM) for 16 h, followed by treatment with LAG-3 shedders (5 or 50 nM) for 6 h. Flow cytometry analysis revealed shedder-mediated degradation was impeded in the presence of the protease inhibitor. This loss of degradation was recapitulated in HeLa cells expressing a cleavage-resistant LAG-3 variant, which was non-responsive to shedder treatment **(b)**. Collectively, these data suggest induced proteolysis is the primary mechanism of action for shedder-mediated LAG-3 degradation.

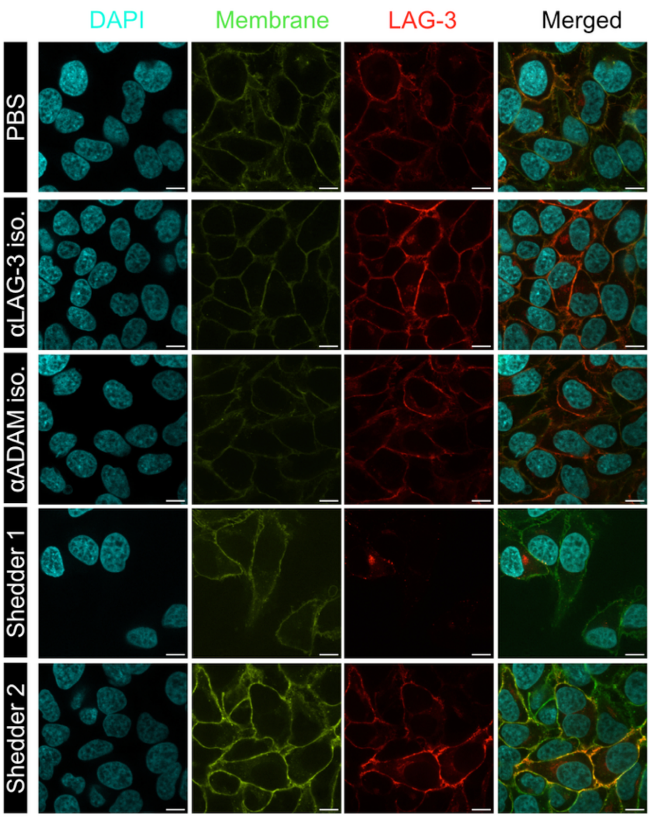


**Figure S5. Visualizing proteolysis-dependent LAG-3 shedding via microscopy.** HeLa cells expressing LAG-3 were pre-treated with ADAM10 inhibitor GI254023X (1 μM) for 16 h, followed by treatment with LAG-3 shedders (50 nM). Representative immunofluorescence images showed that protease inhibition abrogated the activity of shedder 2. In contrast, residual LAG-3 depletion from shedder 1 was still observed ($4^{th}$ panel). This is likely due to the ability of shedder 1 to internalize and shuttle LAG-3 to the lysosome for degradation with when proteolytic cleavage is not available.

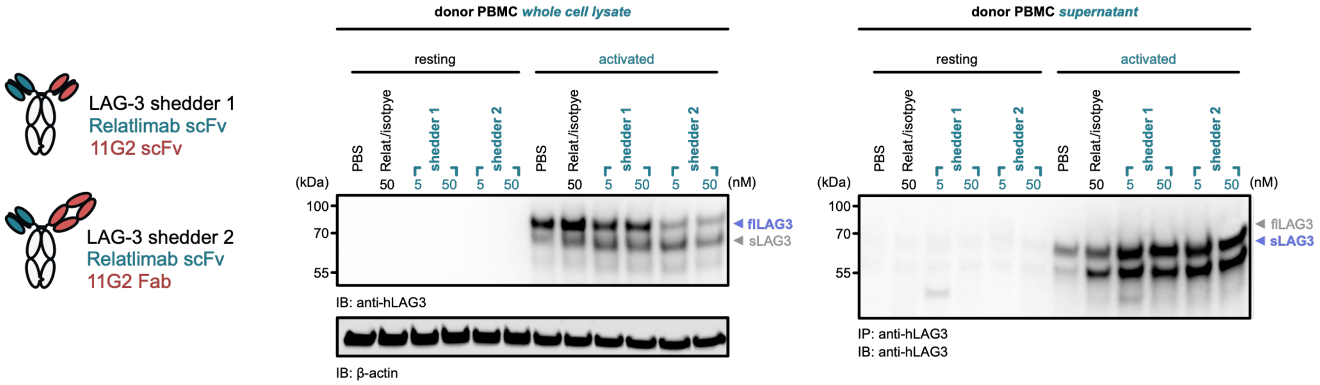


**Figure S6. Induced LAG-3 shedding in human peripheral blood mononuclear cells (PBMCs).** PBMCs were isolated from healthy donors were either in a resting state or activated with immobilized anti-CD3/CD28 (1µg/mL) for 48 h to upregulate LAG-3 expression. These cells were then treated with shedders (5 and 50 nM) and single-arm isotype control (50 nM) for 6 h. Western blot analysis of the whole cell lysate revealed the degree of full-length LAG-3 degradation (left panel), while immunoprecipitation of the culture media revealed the amount of soluble LAG-3 production (right panel). In the absence of activation, no LAG-3 expression or shedding was observed regardless of treatment. Upon activation, both shedders potently degraded LAG-3 from PBMCs with concomitant release of the soluble fragments. These data suggest bispecific-mediated induced shedding is effective in complex, heterogenous environment.

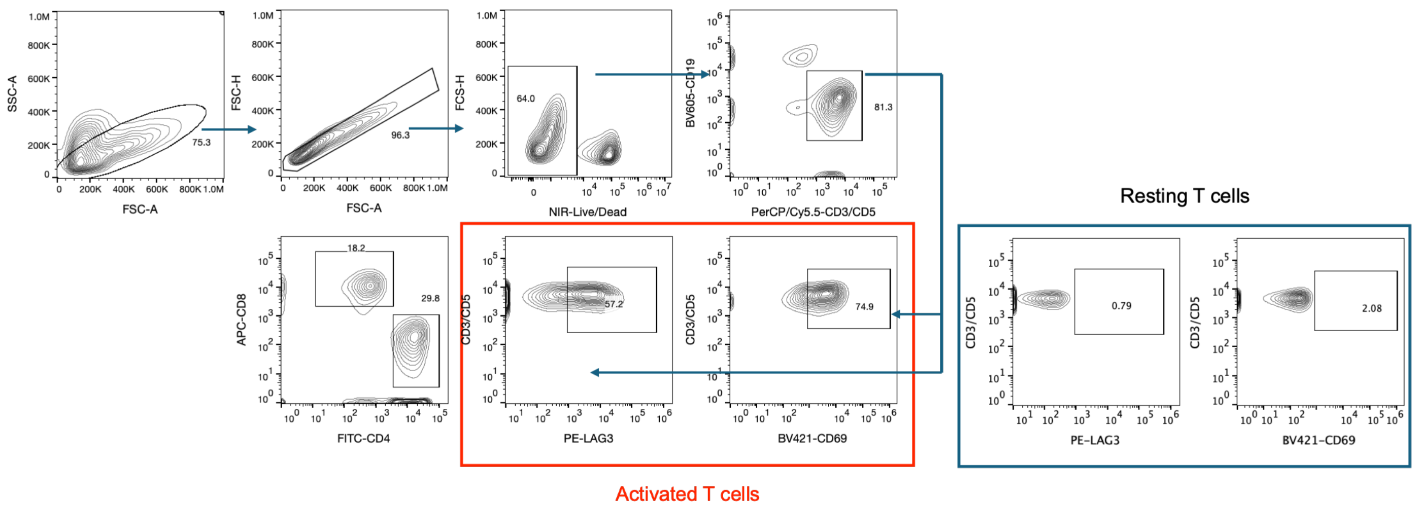


**Figure S7. Gating strategy used to identify LAG-3 shedding on T cells from PBMCs.**

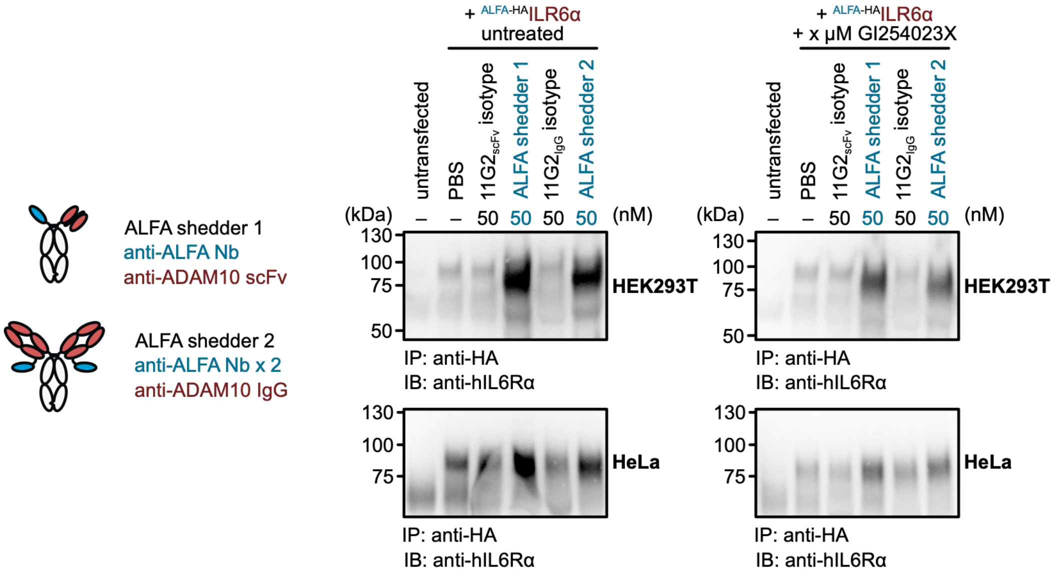


**Figure S8. Induced IL6Rα shedding using the ALFA-tag and universal shedding assay.** HEK293T (top panel) or HeLa (bottom panel) cells were transfected with ALFA-tagged IL6Rα and treated with ALFA shedders or control antibodies (50 nM) for 6 h. Immunoprecipitation of the culture media confirmed enhanced production of soluble IL6Rα (left). This induced IL6Rα shedding necessitates proteolytic function for ADAM10. The release of soluble was drastically reduced when the cells were pre-treated IL6Rα with an ADAM10 inhibitor (right).

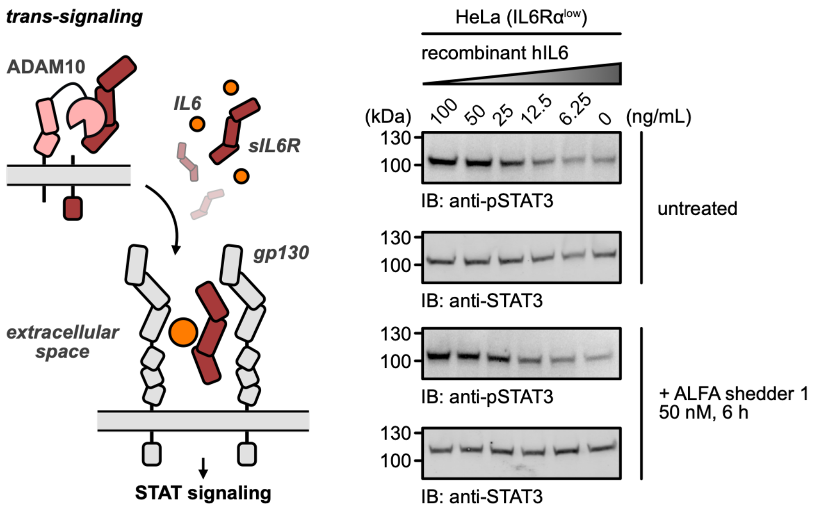


**Figure S9. Induced IL-6Rα shedding using the ALFA-tag and universal shedding assay.** HEK293T cells were transfected ALFA-tagged IL-6Ra and treated with the ALFA shedder (50 nM) to generate soluble IL-6Rα (s IL-6Rα). To access the activity of the shed production, the supernatants from the treated or vehicle-treated cells were mixed with varying concentrations of recombinant IL-6, and transferred to serum-starved HeLa cells. Western blot analysis of the whole-cell lysates (harvested 15 min post-stimulation) revealed that the supernatant from shedder-treated cells (i.e., containing higher level of s-IL-6Rα) comparing to the non-treated controls. These data demonstrate that induced shedding releases a signaling-competent ectodomain capable of facilitating IL-6 trans-signaling.

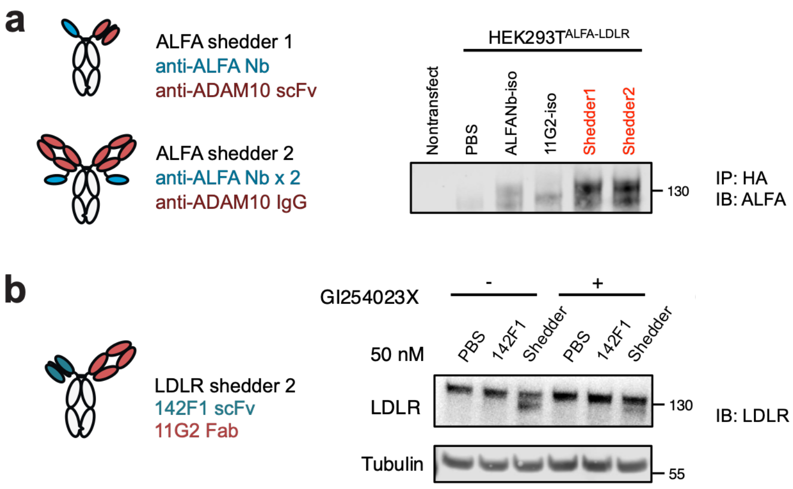


**Figure S10. Induced shedding of LDLR via ALFA- and antibody-based shedders. (a)** HEK293T cells were transfected with ALFA-tagged IL6R$\alpha$ and treated with ALFA shedders or control antibodies (50 nM) for 6 h. Immunoprecipitation of the culture media confirmed the production of soluble LDLR. **(b)** Induced shedding of endogenously expressed LDLR. A bispecific LDLR shedder was engineered using a previously reported anti-LDLR clone 142F1. MDA-MB-231 cells were treated with the LDLR shedder or control antibodies (50 nM) for 6 h. Immunoblotting of the whole cell lysate revealed a second LDLR-species with a lower molecular weight. This product is consistent with the expected ADAM10 cleavage site on LDLR. The generation of this lower molecular weight LDLR species was dependent on ADAM10 activity and was abrogated by pre-treating the cells with GI254023X (1 μM) for 16 h.

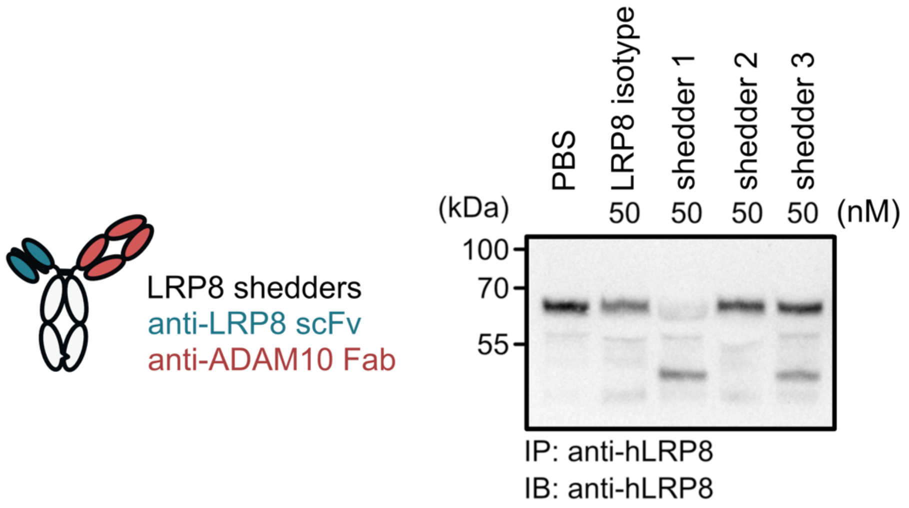


**Figure S11. Induced shedding of LRP8 on SK-N-DZ cells.** SK-N-DZ cells were treated with the LRP8 shedders or control antibodies (50 nM) for 6 h. Immunoprecipitation of culture media from cells treated with the LRP8 shedders reveals distinct cleavage profiles depending on the targeting arm used. While the shedder 3 significantly enhanced both endogenous shedding and the generation of a secondary fragment, the LRP8 shedder 1 produced a unique, lower molecular weight product absent in the PBS control or other shedder treatments. These data suggest that engaging different epitopes on the target can engender novel proteolytic events differ from endogenous cleavage.

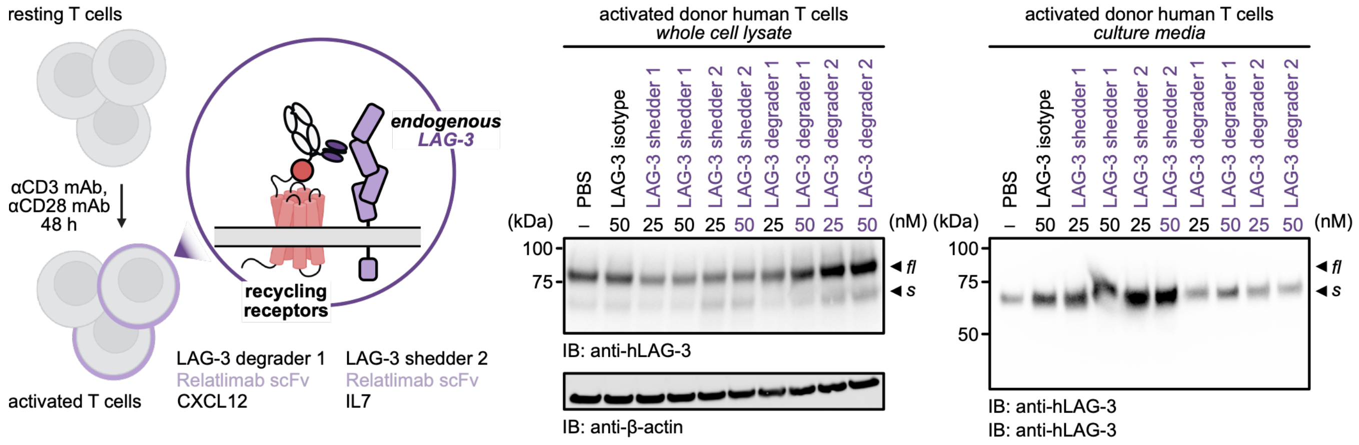


**Figure S12. LAG-3 degradation on primary T cells via bispecific shedders or KineTACs.** To assess the difference between shedding and lysosomal-degradation, we generated two LAG-3 KineTACs using CXCL12 (targeting CXCR7) and IL-7 (targeting IL-7R). T cells were isolated from donor PBMCs and activated with immobilized anti-CD3/CD28 to induced LAG-3 expression. These cells were treated with LAG-3 shedders (50 or 25 nM), KineTACs (50 or 25 nM), and single-arm control antibody (50 nM) for 6 h. Western blot analysis of the whole cell lysate revealed the degree of full-length LAG-3 degradation (left panel), while immunoprecipitation of the culture media revealed the amount of soluble LAG-3 production (right panel). Robust degradation was observed from the LAG-3 shedders and the CXCL12-based LAG-3 KineTAC. At 6 h, both shedders exhibited more potent LAG-3 depletion comparing to the KineTACs. In contrast, soluble LAG-3 was only observed in samples treated with the shedders. These data suggest that induced shedding provides superior degradation kinetics than internalization with the additional benefit of generating functional, soluble fragments.

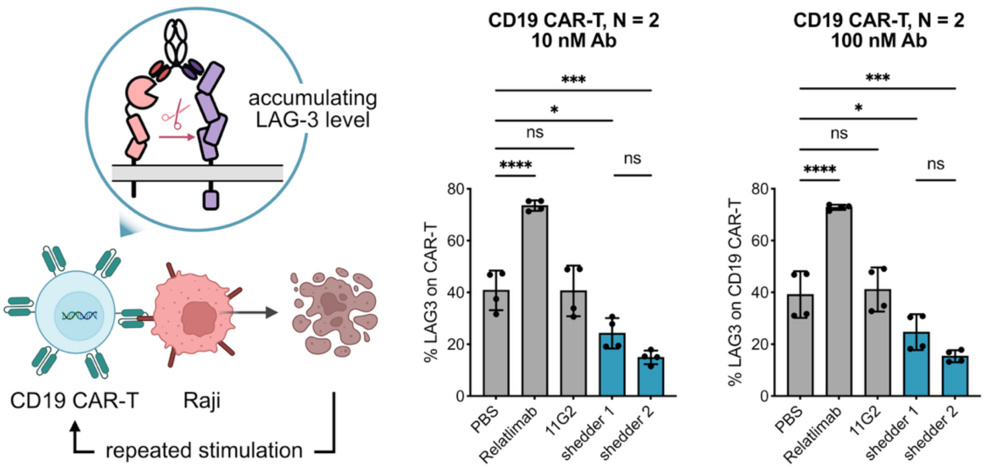


**Figure S13. Induced LAG-3 shedding on CAR-T cells.** A CD19-targeting CAR-T model was used to evaluate LAG-3 shedding in a a therapeutically relevant context. Donor T cells were transduced with CD19-targeting CAR and expanded in vitro with media supplemented anti-CD3/CD28 Dynabeads, IL7, and IL15. Following expansion, CAR-T cells were subjected to 4 rounds of stimulation with $CD19^+$ Raji cells at an effector-to-target (E:T) ratio of 1:1 to induce LAG-3 expression. Next, the CAR-T cells were stimulated one more time with fresh Raji cells in the presence of the LAG-3 shedders, Relatlimab, and 11G2 antibodies (10 nM or 100 nM) and cell surface LAG-3 level was assessed via flow cytometry. Consistent with primary T cell data, Relatlimab upregulated cell surface LAG-3 level on stimulated CAR-T cells. In contrast, LAG-3 shedder 2 potently reduced the amount of LAG-3 on CAR-T cells comparing to non-treated and isotype controls. These data suggest that the bispecific shedders can effectively degrade LAG-3 in a cell therapy context. Percent LAG3 was gated from lymphocyte/singlet/live/GFP+ CAR-T cells. Each sample was tested in biological replicates and error bars represented standard deviation. Statistics were calculated by one-way ANOVA. ns, not significant. *P < 0.05. ***P < 0.001. ****P < 0.0001.
